# Pancreatic cancer cachexia is mediated by PTHrP-driven disruption of adipose *de novo* lipogenesis

**DOI:** 10.1101/2025.06.03.657464

**Authors:** Nikita Bhalerao, Yamini Ogoti, Jessica Peura, Calvin Johnson, Qingbo Chen, Ekaterina D. Korobkina, Faith N. Keller, Maximilian Wengyn, Robert J. Norgard, Claire Shamber, Kelsey Klute, Dominick DiMaio, Karine Sellin, Paul M. Grandgenett, Rui Li, Michael A. Hollingsworth, Adilson Guilherme, Michael P. Czech, Richard Kremer, Lihua Julie Zhu, Jessica B. Spinelli, Emma V. Watson, Marcus Ruscetti, David A. Guertin, Jason R. Pitarresi

## Abstract

Pancreatic cancer patients have the highest rates and most severe forms of cancer cachexia, yet cachexia etiologies remain largely elusive, leading to a lack of effective intervening therapies. Parathyroid hormone-related protein (PTHrP) has been clinically implicated as a putative regulator of cachexia, with serum PTHrP levels correlating with increased weight loss in PDAC patients. Here we show that cachectic PDAC patients have high expression of tumor PTHrP and use a genetically engineered mouse model to functionally demonstrate that loss of PTHrP blocks cachectic wasting, dramatically extending overall survival. The re-expression of PTHrP in lowly cachectic models is sufficient to induce wasting and reduce survival in mice, which is reversed by the conditional deletion of the PTHrP receptor, *Pth1r*, in adipocytes. Mechanistically, tumor-derived PTHrP suppresses *de novo* lipogenesis in adipocytes, leading to a molecular rewiring of adipose depots to promote wasting in the cachectic state. Finally, the pharmacological disruption of the PTHrP-PTH1R signaling axis abrogates wasting, highlighting that a targeted disruption of tumor-adipose crosstalk is an effective means to limit cachexia.

**STATEMENT OF SIGNIFICANCE:** Pancreatic ductal adenocarcinoma (PDAC) is the prototypical cancer type associated with cancer cachexia, a debilitating wasting syndrome marked by adipose tissue loss and muscle atrophy. Herein, we establish that PTHrP is a tumor-derived factor that facilitates cachexia by downregulating *de novo* lipogenesis in adipocytes and that blocking PTHrP is an effective means to limit wasting in preclinical mouse models.

## INTRODUCTION

Pancreatic cancer ranks as the third most fatal cancer in the United States, surpassed only by lung cancer and colorectal cancer [1]. The lack of robust treatment options and early detection opportunities for Pancreatic Ductal Adenocarcinoma (PDAC) patients contribute to a dismal 5-year survival rate of ∼13% [2]. In addition, PDAC patients often present with cancer-associated cachexia, a systemic metabolic syndrome marked by profound adipose tissue and muscle wasting, which impacts body condition and tolerance to available treatment options, thus contributing to reduced survival rates. A striking >75% of PDAC patients suffer from cachexia [3], with nearly one-third of PDAC-associated deaths being attributable to cachexia rather than tumor burden [4]. The perturbed metabolic state in cachectic patients across cancer types has been proposed to be mediated by various tumor-derived factors, such as IL-6 [5], Parathyroid Hormone-Related Protein (PTHrP) [6], TNFα [7], and GDF15 [8, 9], that likely act in concert to alter tumor-host interactions and facilitate wasting.

Clinically, cachexia presents as a multifactorial metabolic syndrome characterized by loss of body weight due to wasting of adipose tissue and muscle mass [10]. Despite patients often presenting with anorexia and anemia, nutritional supplementation does little to curtail weight loss and extend survival [11]. Thus, it is clear from clinical evidence and preclinical models that cachexia is a comorbidity in cancer and that a deeper understanding of the molecular basis of cancer-associated cachexia is required to find new ways to treat cachectic patients and, ultimately, extend survival. Additionally, how tumors communicate with peripheral tissues such as adipose and muscle is largely unexplored and represents an untapped area to apply targeted therapy to block cachectic wasting.

Adipose tissue loss has recently been shown to precede muscle loss in cachectic PDAC patients [12], nominating tumor-adipose crosstalk as a potential point of therapeutic intervention. Additionally, pancreatic cancer patients who exhibit fat only wasting have similarly reduced survival relative to those that exhibit both muscle and fat wasting, demonstrating that loss of adipose is a significant factor affecting survival in PDAC [13]. Under normal physiological conditions, white adipocytes have unilocular lipid droplets and are involved in energy storage and mobilization. During cachexia, some white adipocytes shift their metabolism towards an energy-expending state that resembles thermogenesis [14]. This shift from a primarily energy-storing organ to an energy-consuming organ is a hallmark of adipose tissue wasting in cancer cachexia, yet few tumor-derived factors have been nominated as putative regulators of this crosstalk. Canonically, this metabolic shift has been attributed to an upregulation of adipocyte lipolysis, or the breaking down of fatty acids [15–17]. However, emerging work suggests that decreased *de novo* lipogenesis (DNL) may also contribute to adipose tissue loss in this context [18].

To better understand the molecular etiology of cancer cachexia and associated adipose tissue wasting, we employ a murine model of pancreatic cancer that is accompanied by severe cachexia. We identify PTHrP as a PDAC tumor-derived factor that promotes cancer-associated adipose wasting marked by profound downregulation of *de novo* lipogenesis, and provide pre-clinical evidence to support anti-PTHrP therapy as a cachexia blocking strategy.

## RESULTS

### Tumor cell-specific Pthlh deletion extends overall survival and reduces PDAC-associated cachexia

The conditional expression of *Kras^LSL-G12D^* and *Trp53^LSL-R172H^* were targeted to the murine pancreas using the *Pdx1-Cre* allele, as previously described [19]. The addition of a Rosa26^LSL-YFP^ lineage label allows for the tracking of tumor cells derived from the Pdx1 lineage [20]. The resulting ***<u>K</u>****ras^LSL-G12D^*; *Tr**<u>p</u>**53^LSL-R172H^*; *Pdx1-**<u>C</u>**re*; Rosa26^LSL-**<u>Y</u>**FP^ (KPCY) mice were combined with the *Pthlh^LoxP^*allele (*Pthlh* gene encodes the PTHrP protein) to delete *Pthlh* specifically in tumor cells (Supplemental Figure 1A). Our prior work revealed a genetic linkage between *Kras* and *Pthlh* (these genes are directly adjacent in the mouse and human genome), which precludes the inheritance of homozygous *Pthlh^LoxP/LoxP^* alleles in combination with *Kras^LSL-G12D^* [21]. Nonetheless, the heterozygous deletion of *Pthlh* in KPCY mice (KPCY-Pthlh^HET^) dramatically extended survival compared to KPCY controls, with KPCY-Pthlh^HET^ mice living ∼2.5 months longer (Figure 1A). KPCY-Pthlh^HET^ mice survived a median of 187 days, a ∼68% extension of survival relative to the KPCY median survival of 111 days, which we had previously attributed to reduced metastatic ability [21]. Upon closer examination of the overall body condition of KPCY-Pthlh^HET^ animals, we also observed that they were less cachectic than their KPCY counterparts. To quantify cachexia, body weights were tracked in an age-matched cohort of KPCY and KPCY-Pthlh^HET^ mice in the 15 days prior to sacrifice. KPCY mice underwent significant cachectic wasting over time, while KPCY-Pthlh^HET^ mice, on average, continued to gain or maintain their body mass (Figure 1B). At endpoint, the mouse carcass weights were increased in KPCY-Pthlh^HET^ mice relative to KPCY controls (Figure 1C).

**Figure 1.**
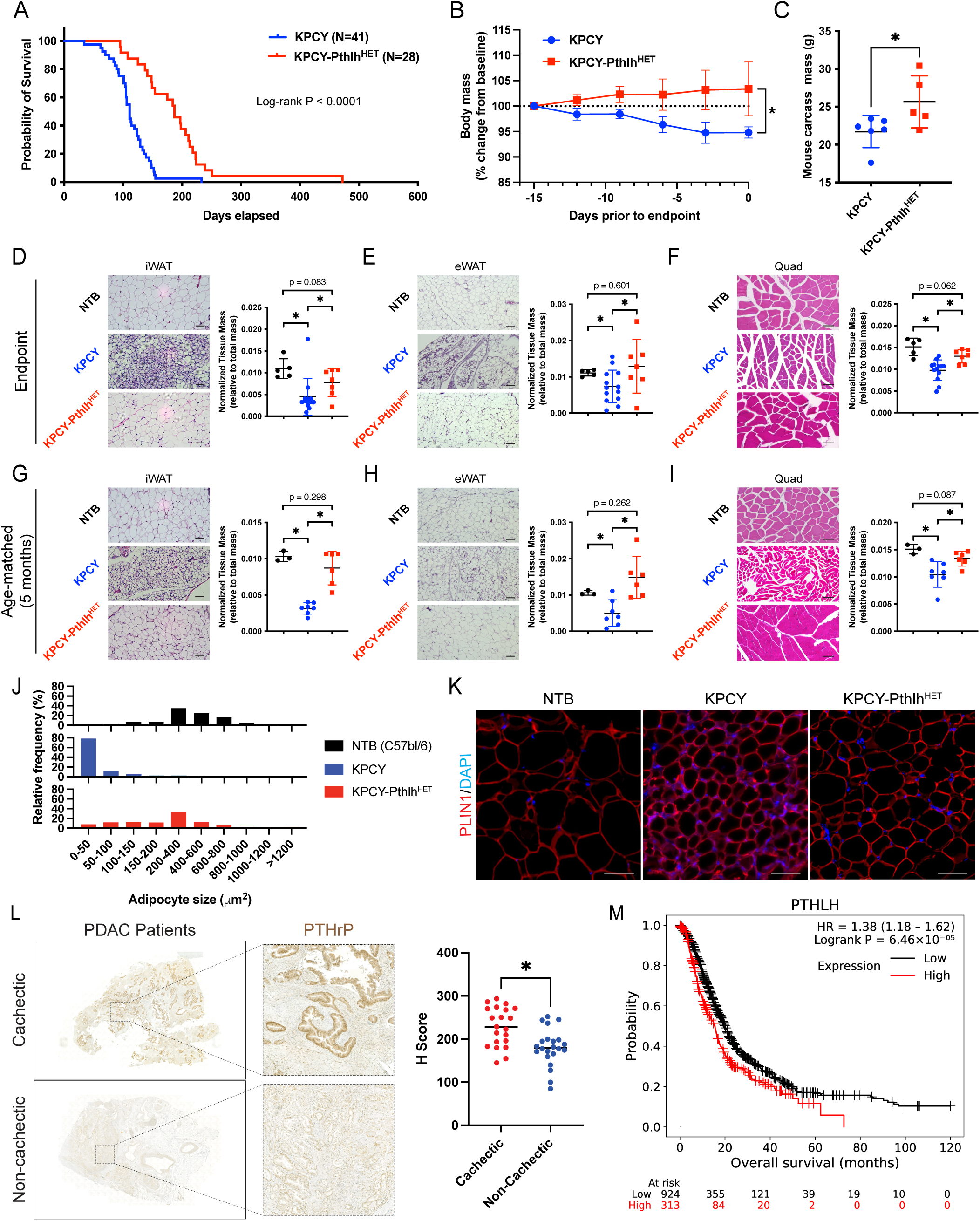
**(A)** Kaplan–Meier overall survival plots of KPCY and KPCY-Pthlh^HET^ mice. Log-rank *P* < 0.0001. **(B)** Percent change from baseline bodyweight in KPCY and KPCY-Pthlh^HET^ mice. **(C)** Carcass weights at endpoint from (B). **(D-F)** Representative H&E images and quantification of normalized tissue masses for iWAT (D), eWAT (E), and quads (F) from non-tumor bearing (NTB), KPCY, and KPCY-Pthlh^HET^ mice at endpoint. Scale bars, 100 μm. **(G-I)** Representative H&E images and quantification of normalized tissue masses for iWAT (G), eWAT (H), and quads (I) from NTB, KPCY, and KPCY-Pthlh^HET^ mice at a set endpoint of 5 months of age. Scale bars, 100 μm. **(J)** Histograms of adipocyte size in iWAT from NTB, KPCY, and KPCY-Pthlh^HET^ mice. **(K)** Immunofluorescence staining of PLIN1 in iWAT from NTB, KPCY, and KPCY-Pthlh^HET^ mice. Scale bars, 50 μm. **(L)** Representative images and H Score quantification of PTHrP IHC in primary tumors from cachectic (N=21) and non-cachectic (N=22) PDAC patients. Scale bars, 100 μm. **(M)** Kaplan-Meier overall survival plots of pancreatic cancer patients stratified by high *PTHLH* (red) or low *PTHLH* (black) expression from the KMPlotter database [59, 60]. Median survival of the high *PTHLH* expressing group is 15.5 months versus 20.0 months for the low *PTHLH* expressing group; Logrank P = 6.46 × 10^−5^. Statistical analysis by Mann– Whitney *U* test (B, C, and D) or Student unpaired *t* test (D, E, F, G, H, I and L), with significance indicated (*, *P* < 0.05). Error bars indicate standard deviation (SD).

To profile more deeply the degree of cachectic wasting in these animals, inguinal white adipose tissue (iWAT), epididymal white adipose tissue (eWAT), and quadricep (quad) muscle were collected across two cohorts; the first set of tissue was collected when mice reached endpoint (Figure 1D-F), and the second cohort at a set endpoint of 5 months (Figure 1G-I). At endpoint, KPCY-Pthlh^HET^ mice showed reduced iWAT, eWAT, and quad wasting (Figure 1D-F). Similarly, in the age-matched 5-month-old cohort, KPCY-Pthlh^HET^ mice showed reduced iWAT, eWAT, and quad wasting (Figure 1G-I). To control for the effects of tumor burden on this analysis, we analyzed the degree of cachexia in a subset of mice from each genotype with matching tumor burden (Supplemental Figure 1B), which revealed that iWAT, eWAT, and quad wasting were still reduced in size-matched KPCY-Pthlh^HET^ mice relative to controls (Supplemental Figure 1C-E). Strikingly, across cohorts, KPCY-Pthlh^HET^ adipose and muscle tissue masses were more similar to those of non-tumor bearing (NTB) animals, suggesting that the loss of tumor cell-derived PTHrP blocked cachectic wasting. Histopathologically, the adipose tissue of KPCY mice was highly cachectic, with marked remodeling of the adipose architecture and a reduction in adipocyte lipid droplet size (Figure 1D-E and 1G-H). In contrast, iWAT and eWAT from KPCY-Pthlh^HET^ mice more closely resembled that of NTB hosts and were primarily composed of large unilocular adipocytes (Figure 1D-E and 1G-H). Muscles from KPCY exhibit reduced fiber size and increased interstitial space, consistent with features of atrophy, while muscles from KPCY-Pthlh^HET^ mice were more similar to NTB animals (Figure 1F and 1I). Cachectic muscle from KPCY mice had increased expression of the E3 ubiquitin ligases *Trim63* (MURF1) and *Fbxo32* (ATROGIN1) that mediate muscle wasting [22], which were lost in KPCY-Pthlh^HET^ mice (Supplemental Figure 1F). To precisely quantify the cachectic burden in adipose tissue, we measured adipocyte size, a proxy for the lipid storing capacity of adipocytes. Adipocyte size was reduced in the iWAT from *KPCY* mice (indicative of adipose wasting), while KPCY-Pthlh^HET^ adipocytes were larger and more similar in size to NTB animals (Figure 1J; Supplemental Figure 1G). Fluorescent staining of iWAT with the adipocyte lipid droplet marker Perilipin-1 (PLIN1) indicated that overall lipid size was reduced in cachectic KPCY mice, which returned to normal in non-cachectic KPCY-Pthlh^HET^ mice (Figure 1K).

Importantly, overall tumor burden did not correlate with the degree of adipose (Supplemental Figure 1H; Pearson R^2^=0.002) or muscle wasting (Supplemental Figure 1I; Pearson R^2^=0.015) across our cohort of KPCY orthotopic and GEMM animals, consistent with prior reports demonstrating that tumor burden does not dictate the degree of cancer-associated cachexia in other models [23, 24]. Indeed, we often observed relatively small tumors (∼500-750mg) that were highly cachectic and much larger tumors (>2 grams) that showed no evidence of cachectic wasting (Supplemental Figure 1H-I). Taken together, the varying degree of PDAC-associated cachexia in genetically identical *KPCY* tumor bearing animals indicates that the ability to induce cachexia is a tumor cell-intrinsic trait, and we have identified highly cachectic and lowly cachectic KPCY tumor cell clones to test putative cachexia-inducing tumor-derived factors such as PTHrP.

Prior work has established that cachectic PDAC patients have higher serum PTHrP relative to non-cachectic patients [25]. We hypothesized, based on our prior work, that PTHrP is coming from PDAC tumor cells [21], and set out to determine if high PTHrP tumor expression corresponds with cachectic burden in a cohort of cachectic and non-cachectic PDAC patients. Indeed, levels of PTHrP in primary tumors from cachectic PDAC patients were increased relative to non-cachectic PDAC patients (Figure 1L), and we have previously demonstrated that increased PTHrP staining in primary tumors correlates with decreased survival of PDAC patients [21]. In agreement with this prior analysis, higher *PTHLH* levels correlate with decreased survival in pancreatic cancer patients from The Cancer Genome Atlas (TCGA) and KMPlotter databases (Figure 1M and Supplemental Figure 1J). PDAC patients with high *PTHLH* levels have a median overall survival of 15.5 months versus 20.0 months in those with low *PTHLH* levels (Figure 1M). Collectively, these data show that PTHrP mediates adipose and muscle wasting cachectic phenotypes in genetically engineered mouse models of pancreatic cancer and that PTHrP is increased in cachectic PDAC patients and correlates reduced with overall survival, setting the stage for mechanistic studies to determine the role of tumor-derived PTHrP in the induction of pancreatic cancer cachexia.

### Pthlh overexpression is sufficient to induce cachexia

The genetic deletion of PTHrP reduced cachexia phenotypes, demonstrating that it is necessary for PDAC-associated cachectic wasting. To determine if PTHrP is sufficient to induce wasting, we took a lowly cachectic KPCY cell line with low basal expression of *Pthlh* (Supplemental Figure 2A) and stably re-expressed *Pthlh* (PTHrP^OE^) via lentiviral-mediated transduction (Supplemental Figure 2B). The resulting KPCY-PTHrP^OE^ tumor cells had a modest, but statistically significant increase in tumor cell proliferation *in vitro* relative to empty vector expressing cells (KPCY-EV; Supplemental Figure 2C), consistent with our prior reports [21]. Upon orthotopic implantation, the re-expression of *Pthlh* significantly reduced overall survival (Figure 2A) and was sufficient to induce a dramatic cachectic phenotype, effectively converting a non-cachectic PDAC cell line into a cachectic one (Figure 2B). Consistent with the induction of cachexia, iWAT architecture was altered upon *Pthlh* overexpression (Figure 2C), and iWAT mass was greatly reduced in KPCY-PTHrP^OE^ tumor bearing animals relative to isogenic KPCY-EV controls (Figure 2D). Adipocyte size was also reduced (Figure 2E; Supplemental Figure 2D) with a concomitant reduction in PLIN1-positive lipid droplet size in iWAT from KPCY-PTHrP^OE^ mice (Figure 2F). Similarly, muscle architecture was perturbed, and muscle mass was reduced in KPCY-PTHrP^OE^ tumor bearing mice (Figure 2G-H). Murf1 (*Trim63*) and Atrogin1 (*Fbxo32*) expression were increased in cachectic muscle from KPCY-PTHrP^OE^ tumor bearing hosts relative to those with KPCY-EV tumors (Supplemental Figure 2E).

**Figure 2.**
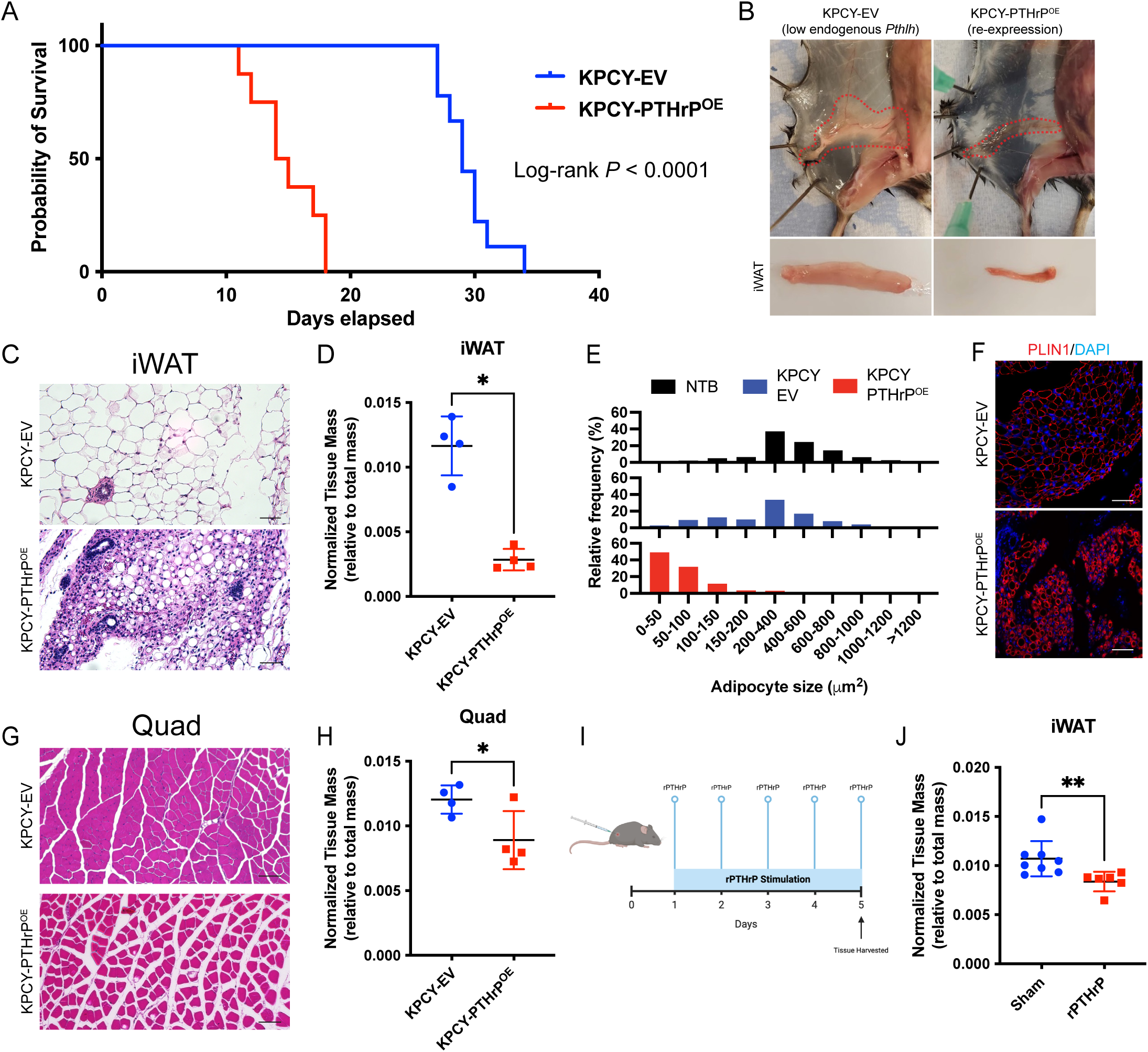
**(A)** Kaplan–Meier overall survival plots of C57BL/6 mice orthotopically implanted with KPCY-PTHrP^EV^ (N=9) or KPCY-PTHrP^OE^ (N=8) cells. Log-rank *P* < 0.0001. **(B)** Representative dissection images of iWAT from KPCY-PTHrP^EV^ or KPCY-PTHrP^OE^ orthotopic tumor-bearing mice. Scale bars, 1 cm. **(C-D)** Representative H&E images and quantification of normalized tissue masses for iWAT from KPCY-PTHrP^EV^ or KPCY-PTHrP^OE^ orthotopic tumor-bearing mice. Scale bars, 100 μm. **(E)** Histograms of adipocyte size in iWAT from KPCY-PTHrP^EV^ or KPCY-PTHrP^OE^ orthotopic tumor-bearing mice. **(F)** Immunofluorescence staining of PLIN1 in iWAT from KPCY-PTHrP^EV^ or KPCY-PTHrP^OE^ orthotopic tumor-bearing mice. Scale bars, 50 μm. **(G-H)** Representative H&E images and quantification of normalized tissue masses for quads from KPCY-PTHrP^EV^ or KPCY-PTHrP^OE^ orthotopic tumor-bearing mice. Scale bars, 100 μm. **(I-J)** Schematic depicting rPTHrP treatment once per day over five days and quantification of normalized tissue masses of iWAT from rPTHrP- or sham-injected C57BL/6 mice. Representative Statistical analysis by Student unpaired *t* test (D and H) with significance indicated (*, *P* < 0.05). Error bars indicate standard deviation (SD).

KPCY-PTHrP^OE^ tumors were larger than KPCY-EV tumors (Supplemental Figure 2F), potentially complicating the interpretation that PTHrP is driving wasting phenotypes. To control for this, we repeated the experiment in an independent cohort where primary tumors were size matched (Supplemental Figure 2G), by harvesting mice at day 18 for KPCY-EV tumors and day 10 for KPCY-PTHrP^OE^ tumors. Despite having less time to induce cachexia, KPCY-PTHrP^OE^ tumor bearing animals showed increased adipose wasting phenotypes in iWAT (Supplemental Figure 2H). At this early time point, quad weights were not significantly different (Supplemental Figure 2I), highlighting that adipose wasting precedes muscle wasting in this model and that PTHrP directly mediates adipose wasting. Finally, to demonstrate that PTHrP acts directly on adipose tissue, we injected non-tumor bearing C57BL/6 animals with recombinant PTHrP (rPTHrP), which was sufficient to decrease iWAT mass after just 5 days of treatment (Figure 2I-J). These results indicate that tumor cell-derived PTHrP is sufficient to induce PDAC-associated cachexia, primarily by facilitating adipose wasting.

### Adipocyte-specific deletion of Pth1r attenuates PDAC-associated cachexia

To decouple tumor growth and energy consumption cachectic phenotypes from direct tumor-adipocyte crosstalk mediated wasting, we set out to disrupt PTHrP signaling by conditionally deleting its cognate receptor, Parathyroid Hormone 1 Receptor (*Pth1r*), in cachectic tissue. Analysis of the Genotype-Tissue Expression (GTEX) database indicated that amongst tissues that undergo wasting in cachexia (adipose, skeletal muscle, heart, etc.), adipose tissue has the highest levels of *PTH1R* expression, with very little to no expression in skeletal muscle (Figure 3A; all tissues shown in Supplemental Figure 3A). To confirm this phenomenon in mice, we isolated RNA from multiple organs of non-tumor bearing C57BL/6 mice and performed qPCR for *Pth1r*, which indicated that adipose tissue depots were the most highly expressing organ that undergoes cachectic wasting (Figure 3B). Of note, *Pth1r* levels are highest in the kidney, consistent with previous studies [26] and likely unrelated to cachectic wasting. Additionally, analysis of a single-cell atlas of human adipose tissue [27] indicated that *PTH1R* is most highly expressed in adipocytes and pre-adipocytes (adipose stem and progenitor cells or ASPCs; Figure 3C), suggesting that adipocyte lineages are the most likely cell in adipose tissue to respond to tumor-derived PTHrP. Similar results were seen in the murine adipose tissue single-cell atlas [27], with *Pth1r* being most highly expressed in adipocytes and pre-adipocytes (Supplemental Figure 3B). Immunofluorescent staining of adipose tissue confirmed that PTH1R is expressed in adipocytes and is increased in cachectic animals (Figure 3D). Collectively, this indicated that PTH1R is highly expressed in adipocytes and is likely to mediate PTHrP-PTH1R tumor to adipose crosstalk.

**Figure 3.**
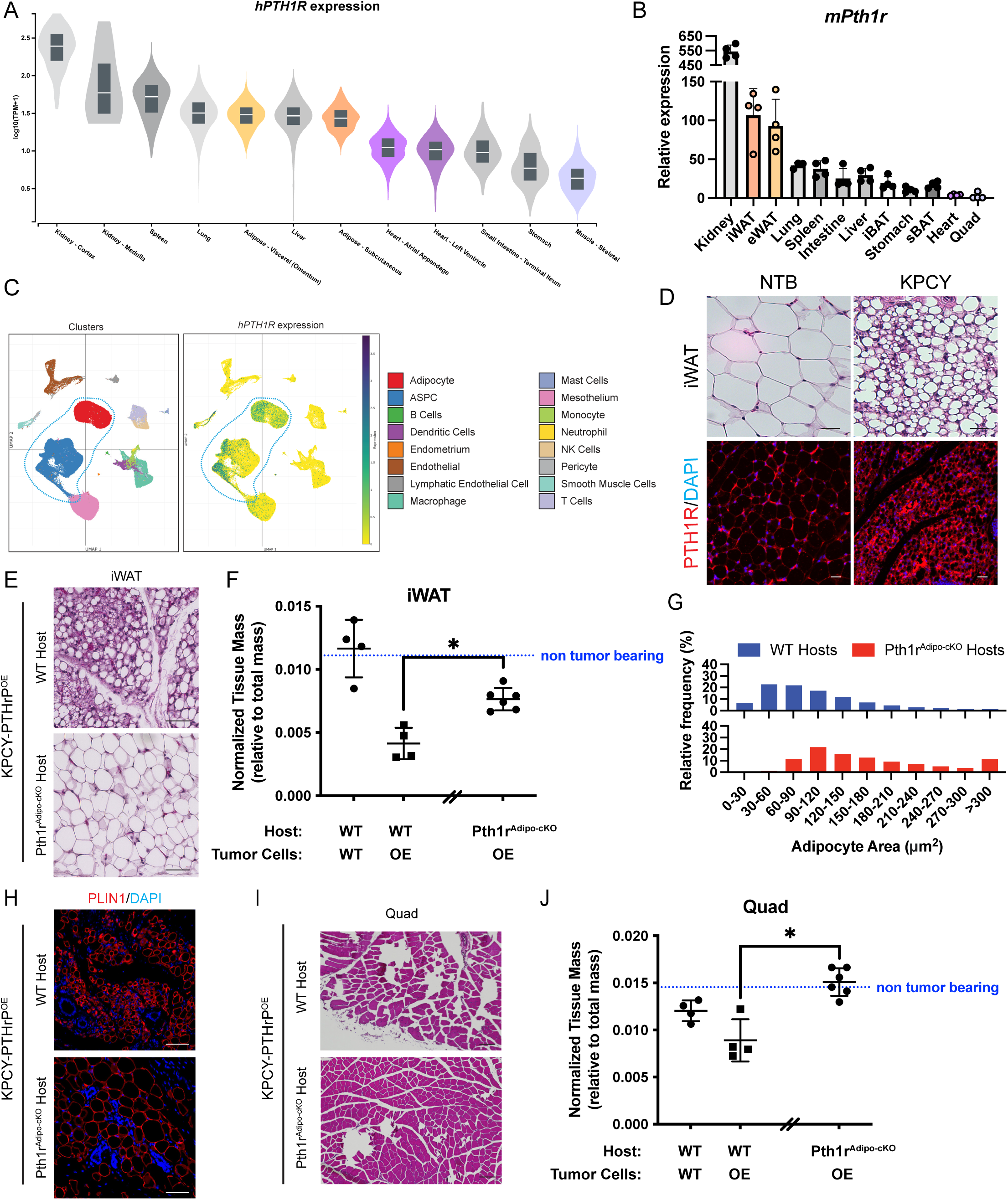
**(A)** Human *PTH1R* expression from the GTEX database in the indicated tissues. Adipose and muscle tissues are highlighted. **(B)** qPCR for murine *Pth1r* expression in the indicated tissues; inguinal Brown Adipose Tissue (iBAT), subcutaneous Brown Adipose Tissue (sBAT). Adipose and muscle tissues are highlighted. **(C)** Single-cell RNA-seq clustering plots (left) and *PTH1R* expression (right) from the Emont *et al*., human adipocyte single cell atlas [27]. **(D)** Immunofluorescence staining of PTH1R in iWAT from non-tumor bearing or KPCY mice. Scale bars, 100 μm. **(E-F)** H&E images and quantification of normalized tissue masses for iWAT from KPCY-PTHrP^WT^ or KPCY-PTHrP^OE^ orthotopic tumor-bearing mice with the indicated host genotypes. Scale bars, 100 μm. **(G)** Histograms of adipocyte size in iWAT from KPCY-PTHrP^OE^ orthotopic tumor-bearing WT or *Pth1r^Adipo-cKO^*host mice. **(H)** Immunofluorescence staining of PLIN1 in iWAT from KPCY-PTHrP^OE^ orthotopic tumor-bearing WT or *Pth1r^Adipo-cKO^*host mice. Scale bars, 50 μm. **(I-J)** H&E images and quantification of normalized tissue masses for quads from KPCY-PTHrP^WT^ or KPCY-PTHrP^OE^ orthotopic tumor-bearing mice with the indicated host genotypes (). Scale bars, 100 μm. Statistical analysis by Student unpaired *t* test (F and J), with significance indicated (*, *P* < 0.05). Error bars indicate standard deviation (SD).

To functionally interrogate the role of the PTHrP-PTH1R signaling axis in cachectic wasting, we deleted *Pth1r* specifically in adipocytes (i.e. *Adiponectin-Cre; Pth1r^LoxP/LoxP^* mice, herein *Pth1r^Adipo-cKO^*). At baseline, there are no differences in adipose tissue (iWAT, eWAT, BAT) weights between *Pth1r^Adipo-cKO^* and WT animals [28]. When implanted with KPCY-PTHrP^OE^ tumors, adipose tissue wasting was reduced in *Pth1r^Adipo-cKO^* mice relative to WT hosts (Figure 3E-F), with the loss of *Pth1r* in adipocytes blocking tumor-induced morphological changes to the adipose tissue (Figure 3F). *Pth1r^Adipo-cKO^*tumor bearing hosts exhibited increased lipid droplet size (Figure 3G-H; Supplemental Figure 3C), indicating that cachectic adipose tissue wasting was reduced when PTHrP-PTH1R signaling in adipocytes is disrupted. Remarkably, blocking PTHrP-PTH1R tumor to adipocyte signaling preserved muscle architecture and reduced wasting (Figure 3I-J). In KPCY-PTHrP^OE^ tumor bearing mice, Murf1 (*Trim63*) and Atrogin1 (*Fbxo32*) expression were increased in cachectic muscle from WT hosts, which was reduced in *Pth1r^Adipo-cKO^* hosts (Supplemental Figure 3D). Thus, tumor cell-secreted PTHrP initiates PTH1R-mediated adipose and muscle wasting.

### PTHrP-mediated adipose tissue wasting downregulates de novo lipogenesis

To determine the downstream mechanism by which PTHrP-PTH1R signaling axis mediates cachectic adipose wasting, we performed bulk RNA-sequencing (RNA-seq) analysis on iWAT from cachectic (KPCY-PTHrP^OE^) and non-cachectic (KPCY-EV) tumor bearing mice (Figure 4A). Cachectic adipose tissue from KPCY-PTHrP^OE^ tumor bearing animals and non-cachectic adipose tissue from KPCY-EV clustered separately by differential gene expression analysis (Supplemental Figure 4A) and Principal Component Analysis (PCA; Supplemental Figure 4B), indicating broad transcriptomic changes between the groups, consistent with the striking morphological changes we observed in the cachectic tissues. The top 50 differentially expressed downregulated genes in KPCY-PTHrP^OE^ tumor bearing hosts relative to KPCY-EV tumor bearing hosts were comprised of known regulators of adipocyte cell biology, including *Fasn*, *Pnpla3*, *Nnat*, and *Orm1* (Supplemental Figure 4C; Supplemental Table 1). The top upregulated genes consisted of immune cell markers such as *Marco*, *Ccr6*, and *Cxcr5* (Supplemental Figure 4C; Supplemental Table 1).

**Figure 4.**
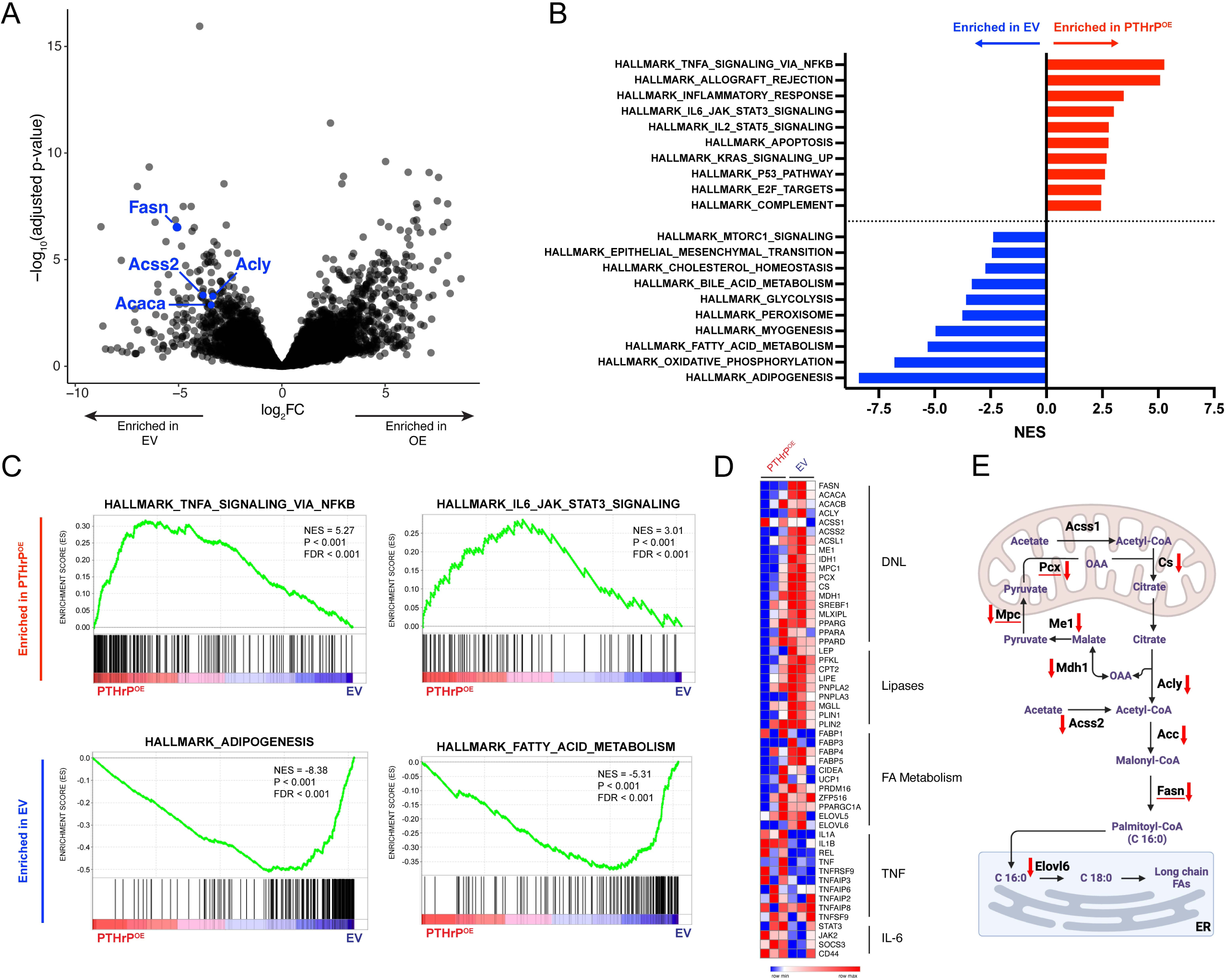
**(A)** Volcano plot of differentially expressed genes from RNA-seq of iWAT from KPCY-PTHrP^WT^ or KPCY-PTHrP^OE^ orthotopic tumor-bearing mice in C57BL/6 hosts; DNL genes highlighted in blue. **(B)** Significantly altered gene set enrichment pathways enriched in iWAT from KPCY-PTHrP^WT^ (blue) or KPCY-PTHrP^OE^ (red) tumor-bearing animals. **(C)** GSEA plots of iWAT from KPCY-PTHrP^WT^ (blue) or KPCY-PTHrP^OE^ (red) tumor-bearing animals. **(D)** Heatmap of altered genes in the DNL, Lipase, Fatty Acid (FA) metabolism, TNF, and IL-6 pathways. **(E)** Schematic representation of the DNL pathway with red arrows indicating all downregulated genes in adipose from KPCY-PTHrP^OE^ tumor-bearing animals relative to KPCY-PTHrP^WT^ tumor-bearing animals. Created in BioRender. Pitarresi, J. (2025) https://BioRender.com/vapcw88

Gene set enrichment analysis (GSEA) indicated that adipose from highly cachectic KPCY-PTHrP^OE^ tumor bearing animals upregulated hallmark mediators of cancer cachexia, such as the TNF-α and IL-6 pathways (Figure 4B-C) [10], consistent with the cachexia-inducing ability of PTHrP [29] . Surprisingly, in addition to these known pro-cachectic pathways, the adipose from KPCY-PTHrP^OE^ tumor bearing animals dramatically downregulated adipogenesis and fatty acid metabolism pathways (Figure 4B-C), suggesting a perturbed regulation of lipid homeostasis in adipocytes during cachexia. Consistent with this pathway analysis, key fatty acid metabolism genes were markedly downregulated in the adipose of highly cachectic animals, including a downregulation of *de novo* lipogenesis (DNL) enzymes (*Acss2*, *Acly*, *Acaca*, and *Fasn*), lipases (*Lep*, *Lipe*, *Pbpla2*, *Plin1/2*), and fatty acid metabolism genes (*FABPs*, *Prdm16*, *Ppargc1a*, *Elovl6*) (Figure 4D). The vast majority of DNL pathway components were downregulated in the adipose from cachectic KPCY-PTHrP^OE^ mice relative to non-cachectic KPCY-EV control mice (Figure 4E; red arrows indicate downregulated genes). Notably, *Fasn* was the most significantly downregulated DNL gene (Figure 4A) and one of the top 15 overall downregulated genes (Supplemental Figure 4C). Thus, tumor-derived PTHrP significantly downregulates adipose DNL to putatively facilitate cachectic wasting.

### Adipocyte specific disruption of de novo lipogenesis sensitizes adipose tissue to wasting

To confirm that DNL pathway components were downregulated in response to PTHrP, we performed qPCR on adipose tissue from PTHrP-overexpressing tumor bearing animals, which showed lower *Acss2*, *Acly*, *Acaca*, and *Fasn* expression (Figure 5A). Western blotting analysis further confirmed the downregulation of DNL enzymes in cachectic iWAT (Figure 5B). Finally, qPCR demonstrated that master transcriptional regulators of DNL, *Mlxipl* (ChREBPɑ and CHREBPβ isoforms) and *Srebf1c* (SREBP), were downregulated in the adipose of PTHrP-overexpressing tumor bearing mice (Figure 5C).

**Figure 5.**
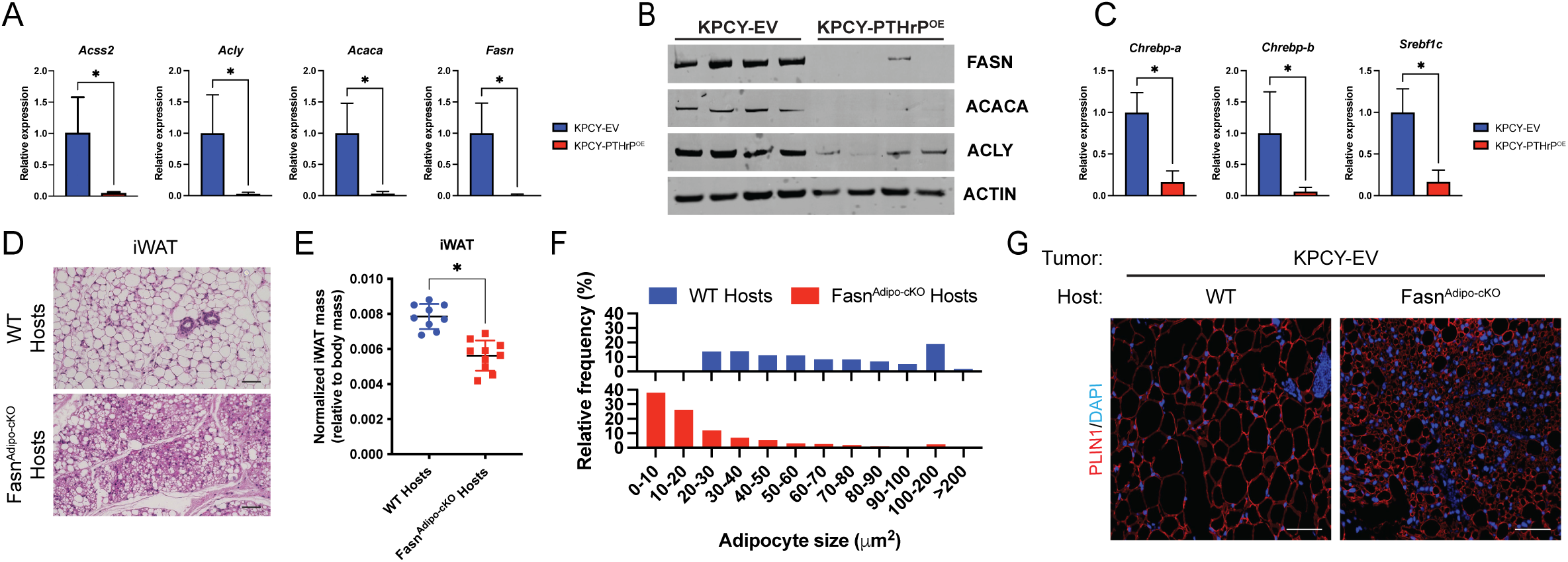
**(A)** qPCR of *Ascc2*, *Acly*, *Acaca*, and *Fasn* in iWAT from KPCY-PTHrP^WT^ or KPCY-PTHrP^OE^ orthotopic tumor-bearing hosts. **(B)** Western blot analysis of FASN, ACACA, ACLY, and ACTIN in iWAT from KPCY-PTHrP^WT^ or KPCY-PTHrP^OE^ orthotopic tumor-bearing hosts. **(C)** qPCR of *Chrebp-a*, *Chrebp-b*, and *Srebf1c* in iWAT from KPCY-PTHrP^WT^ or KPCY-PTHrP^OE^ orthotopic tumor-bearing hosts. **(D-E)** Representative H&E images and quantification of normalized tissue masses for iWAT from KPCY-PTHrP^EV^ orthotopic tumor-bearing C57BL/6 or *Fasn^Adipo-cKO^*hosts. Scale bars, 100 μm. **(F)** Histograms of adipocyte size in iWAT from KPCY-PTHrP^EV^ orthotopic tumor-bearing C57BL/6 or *Fasn^Adipo-cKO^* hosts. **(G)** Immunofluorescence staining of PLIN1 in iWAT from KPCY-PTHrP^EV^ orthotopic tumor-bearing C57BL/6 or *Fasn^Adipo-cKO^* hosts. Scale bars, 50 μm. Statistical analysis by Student unpaired *t* test (A, C, and E), with significance indicated (*, *P* < 0.05). Error bars indicate standard deviation (SD).

To determine the functional role of DNL in PDAC-associated adipose tissue wasting, we implanted the non-cachectic KPCY PDAC line in *Adiponectin-Cre; Fasn^LoxP/LoxP^* mice (herein *Fasn^Adipo-cKO^*; knockout confirmed in iWAT in Supplemental Figure 5A). In this context, the loss of adipocyte *Fasn* resulted in perturbed tissue histology and significantly reduced adipose tissue mass (Figure 5D-E), demonstrating that the disruption of DNL in adipose tissue contributes to cachexia. In addition to worsened adipose tissue wasting, *Fasn^Adipo-cKO^* tumor bearing animals had reduced adipocyte size (Figure 5F; Supplemental Figure 5B) and smaller PLIN1-positive lipid droplets (Figure 5G). Loss of adipose *Fasn* did not alter primary tumor growth (Supplemental Figure 5C) or quad wasting or pathology (Supplemental Figure 5D-E). In non tumor bearing animals, adipocyte specific deletion of *Fasn* had no effect on iWAT or eWAT mass (Supplemental Figure 5F-G), indicating that the loss of *Fasn* leads to cachectic wasting only in the presence of a tumor. Collectively, this data shows a previously unappreciated role for DNL in regulating PDAC-associated adipose tissue wasting.

### Anti-PTHrP therapy blocks PDAC-associated cachexia

No effective therapies currently exist to treat PDAC-associated cachexia. Given our genetic data indicating that the deletion of tumor cell *Pthlh* or adipocyte *Pth1r* reduces cachexia, we set out to determine if anti-PTHrP therapy is an effective means to disrupt PTHrP-PTH1R tumor-adipose crosstalk and block cachectic wasting in PDAC. We have previously demonstrated that monoclonal neutralizing antibodies directed against PTHrP (anti-PTHrP) effectively block PTHrP signaling in PDAC tumor cells, reducing primary tumor burden and metastatic ability [21]. To test the efficacy of anti-PTHrP as a cachexia-blocking therapy in autochthonous KPCY tumors, we enrolled mice when their tumors reached 75 mm^3^, at which point we began anti-PTHrP or anti-IgG antibody treatment. Anti-PTHrP therapy normalized adipose tissue architecture (Figure 6A), reduced adipose wasting (Figure 6B), and increased adipocyte lipid size (Figure 6C; Supplemental Figure 6A) in KPCY mice. Anti-PTHrP treatment of KPCY mice trended towards increased quad mass and normalized morphology (Figure 6D-E). Murf1 (*Trim63*) and Atrogin1 (*Fbxo32*) were mildly induced in the muscles of tumor-bearing animals at this stage, with anti-PTHrP antibody reducing their expression (Supplemental Figure 6B). Similar to the GEMMs, the treatment of orthotopic KPCY tumors (from a cachectic clonal KPCY cell line) with anti-PTHrP therapy showed a similar reduction in cachexia phenotypes, with normalization of the adipose tissue architecture (Figure 6F), reduced adipose tissue wasting (Figure 6G), and increased lipid droplet size (Figure 6H; Supplemental Figure 6C). Additionally, anti-PTHrP treated orthotopic tumor-bearing animals had reduced muscle wasting phenotypes (Figure 6I-J). The muscles of anti-IgG-treated orthotopic mice showed evidence of modest Murf1 (*Trim63*) and Atrogin1 (*Fbxo32*) induction, which was reduced in anti-PTHrP-treated mice (Supplemental Figure 6D). Both KPCY GEMMs and orthotopic tumor bearing animals treated with anti-PTHrP had increased lipid droplet size as visualized by PLIN1 staining of iWAT (Figure 6K-L). Finally, RNA-seq was performed on iWAT from anti-PTHrP (or anti-IgG control) treated orthotopic KPCY tumor bearing animals. Similar to the genetic disruption of PTHrP-PTH1R signaling, anti-PTHrP treatment reduced pro-inflammatory TNF-α signaling and enhanced adipogenesis and fatty acid metabolism signaling pathways in adipose tissue (Figure 6M). Similarly, iWAT from anti-PTHrP treated animals upregulated *Fasn* (Figure 6N) and other DNL enzymes (Supplemental Figure 6E) relative to anti-IgG. Thus, anti-PTHrP monoclonal antibody therapy is an effective means to reduce cachexia in multiple PDAC mouse models, primarily through a reduction in adipose tissue wasting phenotypes by restoring DNL pathway expression.

**Figure 6.**
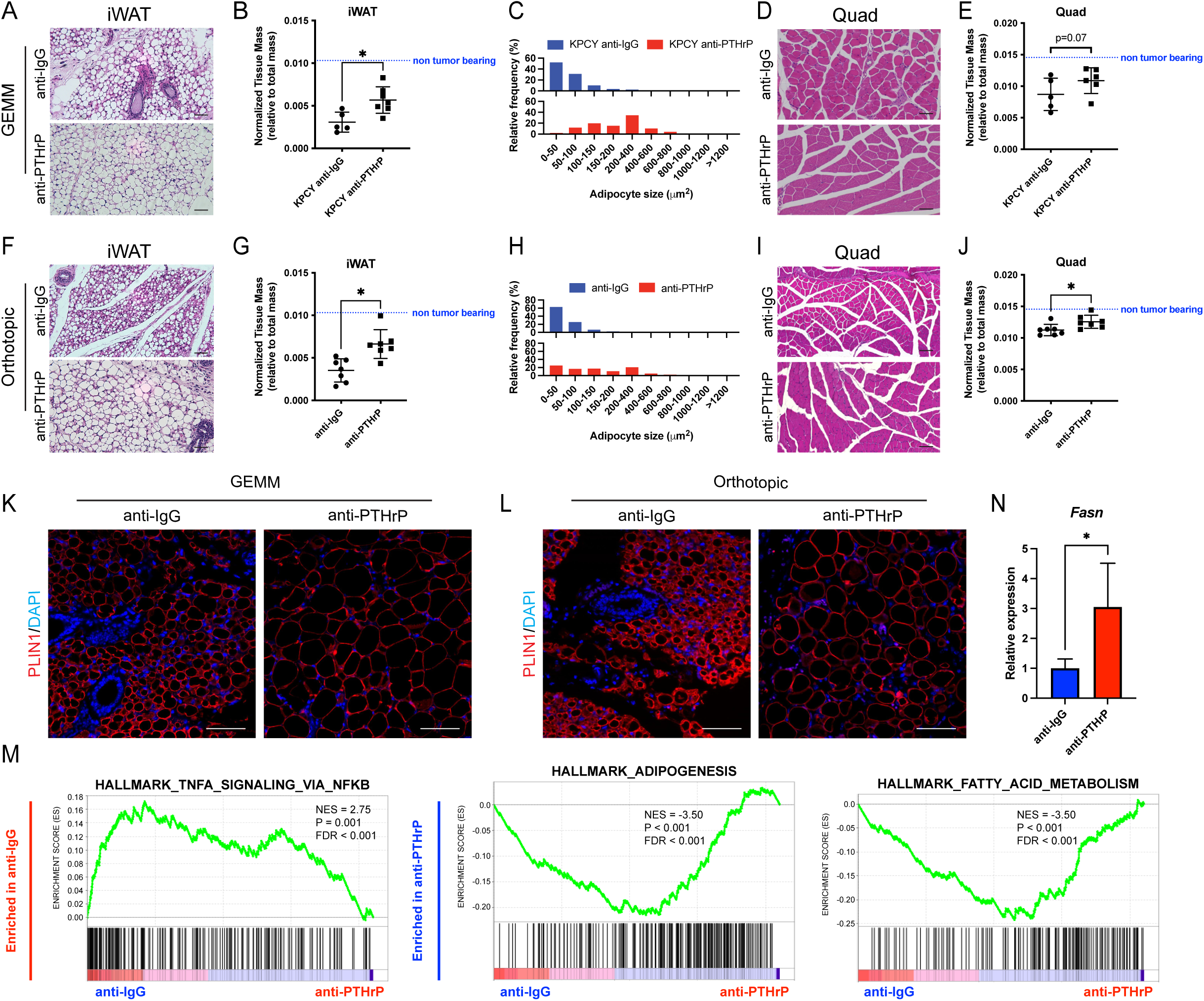
**(A-B)** Representative H&E images and quantification of normalized tissue masses for iWAT from anti-IgG or anti-PTHrP treated KPCY mice. Scale bars, 100 μm. **(C)** Histograms of adipocyte size in iWAT from anti-IgG or anti-PTHrP treated KPCY mice. **(D-E)** Representative H&E images and quantification of normalized tissue masses for quads from anti-IgG or anti-PTHrP treated KPCY mice. Scale bars, 100 μm. **(F-G)** Representative H&E images and quantification of normalized tissue masses for iWAT from anti-IgG or anti-PTHrP treated C57BL/6 mice orthotopically implanted with KPCY cells. Scale bars, 100 μm. **(H)** Histograms of adipocyte size in iWAT from anti-IgG or anti-PTHrP treated C57BL/6 mice orthotopically implanted with KPCY cells. **(I-J)** Representative H&E images and quantification of normalized tissue masses for quads from anti-IgG or anti-PTHrP treated C57BL/6 mice orthotopically implanted with KPCY cells. Scale bars, 100 μm. **(K-L)** Immunofluorescence staining of PLIN1 in iWAT from KPCY mice treated with anti-IgG or anti-PthrP (K) or C57BL/6 mice orthotopically implanted with KPCY cells and treated with anti-IgG or anti-PTHrP (L). Scale bars, 50 μm. **(M)** GSEA plots of iWAT from animals orthotopically implanted with KPCY cells and treated with anti-PTHrP (red) or anti-IgG (blue) tumor-bearing animals. **(N)** qPCR of *Fasn* in iWAT from KPCY orthotopic tumor-bearing hosts treated with anti-IgG or anti-PTHrP therapy. Scale bars, 50 μm. Statistical analysis by Student unpaired *t* test (B, E, G, J, and N), with significance indicated (*, *P* < 0.05). Error bars indicate standard deviation (SD).

## DISCUSSION

Pancreatic cancer remains the lowest 5-year survival rate amongst the common cancers, in large part due to the highest prevalence and severity of cancer cachexia [3]. Cachectic wasting of both the adipose and muscle reduces PDAC patients’ quality of life and overall survival [4, 30], and there are very few treatment options for the clinical management of cancer cachexia. To effectively manage cachexia in the clinic, new therapeutic targets must be identified that directly mediate wasting phenotypes. In our prior work, we identified the oncogenic and pro-metastatic molecule PTHrP as a tumor-derived secreted factor that correlates with decreased survival in human PDAC patients and whose genetic deletion or pharmacological inhibition in PDAC mouse models greatly enhances survival [21]. Herein, we expand upon these initial findings to show that PTHrP contributes to PDAC-associated adipose tissue wasting in cachexia (Figure 7). Thus, targeting PTHrP therapeutically may be a multi-pronged attack to simultaneously block cancer cachexia, tumor growth, and metastasis. We show that cachectic patients have higher tumor PTHrP expression relative to non-cachectic patients, and that the genetic deletion or pharmacological inhibition of PTHrP reduces cachectic wasting in mouse models of PDAC-associated cancer cachexia. Induced expression of *Pthlh* in a lowly cachectic PDAC tumor cell line was sufficient to induce adipose and muscle wasting, ultimately leading to reduced overall survival in mice. Disruption of PTHrP-PTH1R tumor to adipose crosstalk via adipocyte-specific conditional deletion of *Pth1r* reduced adipose and muscle wasting, demonstrating that direct tumor cell to adipocyte signaling networks mediate adipose wasting that ultimately fuels muscle wasting. Mechanistically, we found that cachectic adipose tissues lose core DNL signaling components while concomitantly upregulating pro-cachectic factors such as TNF-α and IL-6. Finally, the adipose-specific deletion of *Fasn* sensitized animals towards adipose wasting, demonstrating that loss of DNL signaling contributes to cachexia. As such, our results reveal a previously unappreciated PTHrP-PTH1R-DNL signaling axis that drives cancer-associated cachexia and provides a promising target to ameliorate cachectic wasting in PDAC patients.

**Figure 7.**
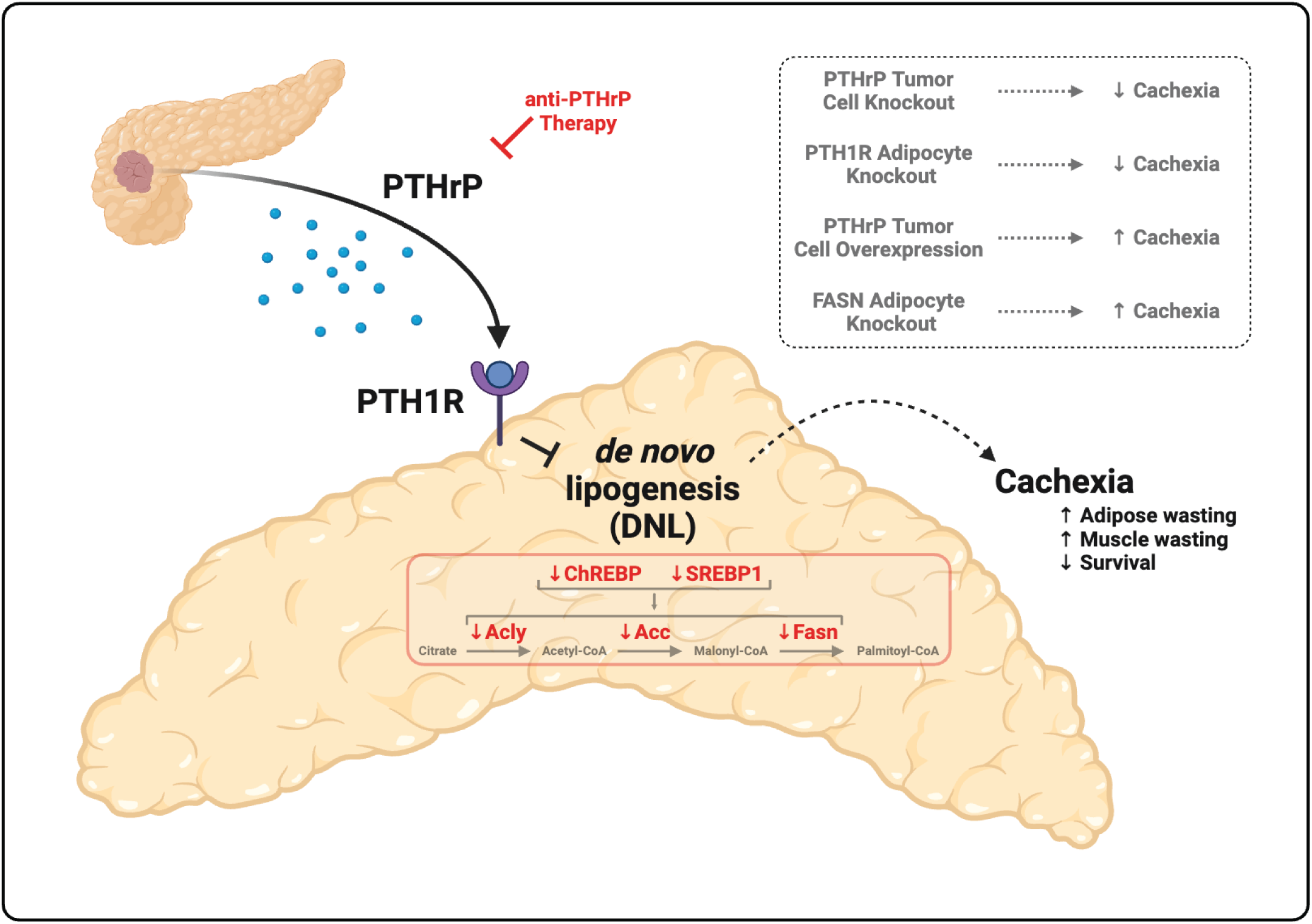
Schematic of PTHrP-induced adipose tissue wasting. Tumor-derived PTHrP binds to PTH1R on adipocytes, which downregulates *de novo* lipogenesis in adipose tissue to facilitate wasting. Created in BioRender. Pitarresi, J. (2025) https://BioRender.com/wbw5raq

Cancer cachexia involves complex multi-organ metabolic crosstalk [10], with tumor cells sending and receiving multiple inputs from systemic paracrine/endocrine signaling networks. Several mechanisms have been identified to date to indicate how tumor cells communicate with the muscle to facilitate muscle wasting and catabolism, but few studies have identified mediators of direct tumor to adipocyte crosstalk as drivers of adipose tissue wasting and subsequent muscle wasting [31]. This is important clinically, as adipose tissue wasting precedes muscle wasting in PDAC patients and can occur up to 1.5 years prior to diagnosis [12], and patients who exhibit fat only wasting have significantly reduced survival [13]. Similar phenomena have been observed in other cancers as well [16, 32, 33]. Our adipocyte-specific deletion of *Pth1r* interestingly halted both adipose wasting and muscle wasting, suggesting that adipose wasting in this model may fuel muscle wasting. Future studies will seek to identify and functionally characterize how these adipose tissue-secreted signaling molecules, called “adipokines”, mediate muscle wasting, either through direct catabolism or by altering the metabolic status of the host.

Adipose tissue wasting during cancer cachexia has primarily been attributed to the induction of a UCP1-driven thermogenic program that promotes adipocyte browning [34]. Indeed, PTHrP is known to trigger adipose tissue browning in the Lewis Lung Carcinoma (LLC) model of cancer cachexia [6], and we find a modest induction of adipose UCP1 in our PDAC-associated cachexia models (Figure 4D). However, we found a more profound downregulation of *de novo* lipogenesis in our system that suppresses lipid production in adipocytes and contributes to cachectic wasting. Thus, harnessing new therapies to block or reverse the loss of DNL hold promise as a means to reverse cachectic wasting of adipose tissue depots. One such approach that shows promise in preclinical models is the Peroxisome proliferator-activated receptor alpha (PPARα) agonist fenofibrate, which has been shown to rescue reduced *Acly*, *Fasn*, *Acsl1*, and *Acsl5* in hepatic *de novo* lipogenesis, thus restoring lipid homeostasis in the liver and improving overall survival [18]. Importantly, our data indicate that PPARα (encoded by *Ppara*) is still expressed, albeit at slightly lower levels, in the adipose of PTHrP tumor bearing animals, and thus its reactivation is an attractive therapeutic target to pursue in future studies.

Prior correlative studies in a small cohort consisting of cachectic (weight losing) and non-cachectic (weight stable) patients suggested that enhanced adipocyte lipolysis, but *not* reduced lipogenesis was involved in adipose tissue loss in cancer cachexia [35]. Provocatively, other work has demonstrated that whole body lipolytic rate in cancer patients is not different from healthy controls, suggesting that the loss of body fat in cancer cachexia is due to reduced lipogenesis rather than augmented lipolysis [36]. The latter is supported by clinical data to show that lipogenic enzymes such as FASN are reduced in adipose from cachectic patients [37, 38], consistent with our experiments showing that PTHrP mediates loss of DNL in cachectic adipose tissue. Nonetheless, subsequent functional studies in the field have primarily focused on the role of enhanced lipolysis, and not reduced DNL, in mediating adipose tissue loss in cachexia. This work has confirmed that lipases are enriched in the cachectic state, which can be blocked through the inhibition of key lipases such as adipose triglyceride lipase (ATGL) or hormone-sensitive lipase (HSL) to limit adipose wasting [16]. Surprisingly, in PTHrP-driven adipose wasting, the expression of most cachexia-associated lipases, such as ATGL (*Pnpla2*), HSL (*Lipe*), and Monoglyceride Lipase (*Mgll*), were unchanged or even reduced in the cachectic state, suggesting that they are not driving the adipose tissue wasting phenotype in this model or that their regulation occurs post-transcriptionally. Rather, our work demonstrates that the loss of *de novo* lipogenesis in adipocytes contributes to adipose tissue wasting. Consistent with this working model, prior work has demonstrated that the decrease in adipose tissue mass is due to reduced adipocyte size and not from a reduction in the total number of adipocytes (i.e. adipocyte death) [35, 39]. It is likely that the balance between adipose tissue lipolysis and lipogenesis is altered during progressive cachexia-associated adipose wasting, and thus different models may find varying functional roles for these enzymes during different stages of cachexia progression.

PDAC patients with higher serum levels of PTHrP lose significantly more weight, have reduced handgrip strength, and score lower on the Karnofsky Performance Status (KPS) scale relative to those with lower serum PTHrP, indicative of enhanced cachectic burden [25]. Additionally, whole body fat oxidation is increased in PDAC patients with high PTHrP, suggesting a perturbed homeostasis between lipolysis and lipogenesis [25], in agreement with our preclinical experiments in mice. Critically, in our models, disrupting direct tumor-to-adipocyte crosstalk by deleting *Pth1r* on adipocytes was sufficient to block cachectic wasting. This result, combined with our prior work showing that PTHrP facilitates tumor growth and metastasis, demonstrates that blocking PTHrP-PTH1R signaling axis in preclinical models is a potent means to block many of the confounding factors in PDAC that lead to its dismal overall survival rate. Interestingly, PTHrP/*PTHLH* is a squamous identity gene and is highly expressed in many squamous carcinomas, including the squamous/basal-like subtype of PDAC [21], lung squamous cell carcinoma [40], head and neck squamous cell carcinoma [41], oral squamous cell carcinoma [42], and cervical squamous cell carcinoma [43]. Many of these squamous cancers also have high levels of cancer-associated cachexia, suggesting that targeting PTHrP in other cancers to block cachectic wasting warrants further preclinical testing. In a recent pan-cancer analysis, patients with high serum PTHrP levels lost ∼7% of their body weight independent of tumor burden, while those without PTHrP in their serum lost only ∼1% [44], highlighting that PTHrP may also drive wasting in other cancers. Notably, the Lewis lung cancer model of cancer cachexia was similarly responsive to anti-PTHrP therapy [6], perhaps pointing towards a more broadly applicable mechanism of PTHrP-induced cachectic wasting across cancer types.

## METHODS

### Mice

*Kras^LSL-G12D^*; *Trp53^LSL-R172H^*; *Pdx1-Cre*; *Rosa26^LSL-YFP^* (KPCY) and *Kras^LSL-G12D^*; *Trp53^LSL-R172H^*; *Pdx1-Cre*; *Rosa26^LSL-YFP^; Pthlh^Loxp/wt^* (KPCY-Pthlh^Het^) mice have been described previously [20, 21]. *Pthlh^LoxP^* mice were a generous gift from Dr. Andrew Karaplis (McGill University) [45]. *Pth1r^LoxP^* mice were a generous gift from Dr. Henry Kronenberg (Harvard University) [46]. Fasn^LoxP^ mice were generously provided by Dr. Clay Semenkovich (Washington University at St. Louis) [47]. Adiponectin-Cre and wild-type C57BL/6J mice were purchased from the Jackson Laboratory (Stock # 028020 and #000664, respectively). *Fasn^LoxP^* or *Pthlh^LoxP^*mice were crossed with Adiponectin-Cre mice to generate *Adiponectin-Cre; Fasn^LoxP/LoxP^* (Fasn^Adipo-cKO^) and *Adiponectin-Cre; Pth1r^LoxP/LoxP^* (Pth1r^Adipo-cKO^) lines, respectively. For survival analysis, mice were palpated for tumors and examined for evidence of morbidity twice weekly and sacrificed when moribund. Mice treated with monoclonal neutralizing antibody against PTHrP (anti-PTHrP; generously provided by Dr. Richard Kremer, McGill University; 200μg/injection) or anti-IgG control were injected three times per week every 48 hours for the indicated time. Mice were treated with rPTHrP (1mg/kg; Bachem T-4512) for 5 consecutive days, and tissues were harvested within 1 hour of the final dosing. All vertebrate animals were maintained, and experiments were conducted in compliance with the NIH guidelines for animal research and approved by the University of Massachusetts Chan Medical School Institutional Animal Care and Use Committee.

### Genotyping

Mice were genotyped by automated genotyping services via TransnetYX for the following alleles: *Kras^LSL-G12D^*, *Trp53^LSL-R172H^*, *Pdx1-Cre*, *Rosa26^LSL-YFP^, Pth1r*, *Adiponectin-Cre*, and *Pthlh^Loxp^*. For Fasn^Adipo-cKO^ mice, conventional PCR genotyping reactions were performed. Briefly, tails were cut and immersed in lysis buffer (40% NaOH + 0.5M EDTA pH 8) for one hour and incubated at 95°C, followed by addition of neutralization buffer (40mM Tris-HCl). PCR reactions were carried out with the following primers:

*Fasn forward: 5’ - GGA TAG CTG TGT AGT GTA ACC AT -3’*

*Fasn reverse: 5’- GGT CAT CGT GAT AAC CAC ACA T -3’*

*Adipo-Cre transgene forward: 5’- ACG GAC AGA AGC ATT TTC CA -3’*

*Adipo-Cre transgene reverse: 5’- GGA TGT GCC ATG TGA GTC TG - 3’*

*Adipo-Cre internal control forward: 5’- CTA GGC CAC AGA ATT GAA AGA TCT -3’*

*Adipo-Cre internal control reverse: 5’- GTA GGT GGA AAT TCT AGC ATC ATC C -3’*

### Orthotopic injection

Orthotopic injection was performed as previously described [48]. Briefly, C57BL/6 mice were anesthetized using isoflurane. A small incision was made on the upper left quadrant of the mouse, and the pancreas was exteriorized on a sterile field. 1.0 × 10^5^ cells were injected into the pancreas in 100ul sterile PBS via an insulin syringe. A cotton swab was held over the site of injection. After injection, the pancreas was placed back into the peritoneal cavity and the incision was sutured with 4-0 Vicryl sutures. The mice received meloxicam post-surgery and were monitored daily following surgery to assess post-surgery recovery. For each experiment, the sex of mice was matched to the sex of the KPCY cell line.

### Immunohistochemistry and immunofluorescence staining

For staining of mouse tissues, zinc formalin-fixed paraffin embedded (FFPE) tissues were dewaxed by baking in the oven at 60°C for an hour. Following dewaxing, tissues were rehydrated by serial immersions in xylene-substituent histoclear and ethanol gradients. Antigen retrieval was performed using citric acid-based antigen retrieval buffer (Vector laboratories H-3300). Plin1 (Cell signaling technologies 9349), PTHrP (Sigma AV33885), and PTH1R (Abcam ab189924) primary antibodies were used at 1:1000 dilution overnight at 4°C. Tissues were incubated with either HRP-tagged secondary antibody (Vector laboratories MP-7401) or donkey anti-rabbit Alexa fluor 594 (Fisher Scientific Cat #A21207) at room temperature for 1 hour and followed with hematoxylin counterstain or mounted with DAPI mounting media. Images were acquired on a Leica Thunder Imager or EVOS M5000.

### Human specimens

Primary PDAC patient tumor samples and associated omental adipose tissues were obtained from decedents who had previously been diagnosed with pancreatic ductal adenocarcinoma from the University of Nebraska Medical Center’s Tissue Bank through the Rapid Autopsy Program (RAP) in compliance with IRB 091-01. Non-cancer tissues are collected in a manner similar to RAP specimens through the UNMC Normal Organ Recovery (NORs) Program in association with LiveOn Nebraska. To ensure specimen quality, organs were harvested within three hours post-mortem and the specimens flash frozen in liquid nitrogen or placed in formalin for immediate fixation. Sections were cut from paraffin blocks of formalin fixed tissue into 4-micron thick sections and mounted on charged slides. Pancreatic cancer patients with a known history of body weight loss, as confirmed by the pathologist, were binned into cachectic and non-cachectic cohorts. The primary tumors were processed and stained for PTHrP immunohistochemistry as described above. The slides were scanned by the UMass Chan SCOPE core facility using a SL Slide Scanner, followed by PTHrP H Score quantification using QuPath software.

### Cell lines and culture conditions

Murine PDAC cell lines were derived from KPCY mice that were backcrossed onto the C57BL/6J strain [49]. KPCY-PTHrP^EV^ and KPCY-PTHrP^OE^ cells were described previously [21]. All cell lines were maintained in Dulbecco’s modified eagle medium (DMEM) with 10% fetal bovine serum (FBS) and 1% penicillin/streptomycin (P/S) at 5% CO_2_ at 37°C. Cell lines were tested regularly for *Mycoplasma*.

### Cell proliferation assay

The WST-1 cell proliferation assay (ab65475, Abcam) was performed per the manufacturer’s instructions. Briefly, 2.0 × 10^3^ cells were plated on 96-well plates and grown in DMEM supplemented with 10% FBS and 1% P/S. Fresh media containing 10% WST solution was incubated on the cells for 2 hours at 37°C. After shaking the plate, the absorbance was read at 420 to 480 nm with a reference wavelength of 650 nm. Final absorbance was obtained by subtracting the 650-nm reference wavelength from the 420- to 480-nm read.

### Western blot analysis

Inguinal white adipose tissues were snap-frozen in liquid nitrogen immediately after retrieving from orthotopic and GEM mice. The iWAT’s were transferred to 2ml Eppendorf tubes with pre-cooled metal bead and homogenized using Cryomill (Retsch) for 3 minutes to get a fine powder. The powderized iWAT’s were then lysed in RIPA buffer containing protease and phosphatase inhibitors (Sigma 04693124001 and Roche 04906837001). The samples were centrifuged at 15,000 g for 15 min at 4°C, and the supernatant was collected for western blotting. Lysates were run on NuPAGE gels (Thermo Fisher NP0336BOX), transferred to PVDF membranes, and stained with primary antibodies: Fasn (Cell signaling technologies 3180), Acly (Proteintech 15421-1-1 AP), Acaca (Cell signaling technologies 3662) and beta-actin (Sigma A5316). Membranes were washed and stained with secondary antibodies: donkey anti-mouse IgG IRDye 800 (LICORbio 926-32212) or donkey anti-rabbit IgG IRDye 800 (LICORbio 926-32213) or donkey anti-rabbit IgG IRDye 680 (LICORbio 926-68073) (1:15,000), washed, and imaged on a LI-COR Odyssey imaging system.

### RNA extraction and qPCR

RNA was extracted from cryomilled adipose tissues using QIAzol/chloroform and Qiagen RNeasy kit (Cat #74106) following the manufacturer’s protocol. For extracting RNA from the iWAT, powderized tissues were resuspended in 1ml QIAzol per sample and lysed using TissueLyser II (Qiagen Cat#) and pre-cooled metal beads. The lysed sample was mixed with chloroform and mixed vigorously, followed by centrifugation at 4°C for 10 minutes at 10,000g. The upper aqueous phase was mixed with 70% ethanol in 1:1 ratio and transferred to the RNA spin columns from the Qiagen RNeasy kit and spun for 20 seconds at 9000g. All the remaining steps were followed as per manufacturer’s guidelines. Extracted RNA was quantified using Nanodrop and cDNA was prepared using 100ng-1000ng RNA using the New England Biolabs LunaScript RT supermix (Cat# M3010L). Quantitative PCR was setup using Luna Universal qPCR Master Mix (Cat #M3003E). The following primers were used:

*Pth1r forward: 5’-TCT GCA ATG GTG AGG TGC AG - 3’*

*Pth1r reverse: 5’- GCT ACT CCC ACT TCG TGC TT - 3’*

*Fasn forward: 5’- GGA GGT GGT GAT AGC CGG TAT - 3’*

*Fasn reverse: 5’- TGG GTA ATC CAT AGA GCC CAG - 3’*

*Acly forward: 5’- ACC CTT TCA CTG GGG ATC ACA -3’*

*Acly reverse: 5’- GAC AGG GAT CAG GTA TTC CTT G - 3’*

*Acaca forward: 5’- ATT GTG GCT CAA ACT GCA GGT - 3’*

*Acaca reverse: 5’- GCC AAT CCA CTC GAA GAC CA -3’*

*Murf1 forward: 5’- GGA CGG AAA TGC TAT GGA GAA CC CC-3’*

*Murf1 reverse: 5’- GAT GGC TGT TTC CAC AAG CTT GG - 3’*

*Atrogin1 forward: 5’- CCT CAG CAG TTA CTG CAA CAA GGA -3’*

*Atrogin 1 reverse: 5’- CAT CTT CTT CCA ATC CAG CTG C - 3’*

*Chrebp-a forward: 5’- CGA CAC TCA CCC ACC TCT TC -3’*

*Chrebp-a reverse: 5’ - TTG TTC AGC CGG ATC TTG TC - 3’*

*Chrebp-b forward: 5’ - TCT GCA GAT CGC GTG GAG -3’*

*Chrebp-b reverse: 5’- CTT GTC CCG GCA TAG CAA C -3’*

*Srebf1c forward: 5’- AAG CAA ATC ACT GAA GGA CCT GG - 3’*

*Srebf1c reverse: 5’- AAA GAC AAG CTA CTC TGG GAG -3’*

*18S RNA forward: 5’- CGA ACG TCT GCC CTA TCA ACT T - 3’*

*18S RNA reverse: 5’- CCG GAA TCG AAC CCT GAT T - 3’*

### RNA-seq library prep and GSEA

Frozen iWAT tissues were cryomilled and RNA was isolated using Qiazol and Qiagen RNAeasy mini kits. A quantity of 500ng total RNA was used for poly(A) mRNA purification using the NEBNext poly(A) mRNA magnetic isolation module (E7490L). Libraries were prepared using the NEBNext Ultra II Directional Library Prep Kit for Illumina (E7760L) as per the manufacturer’s recommendations and were barcoded using NEBNext Multiplex Oligos for Illumina (Dual Index Primers set 2 E7780S) during PCR amplification. Final libraries were quantified using the NEBNext Library Quant Kit for Illumina (E7630L). Sequencing was performed using an Illumina NextSeq2000 with 115 bp single-end reads, plus 8 cycles for barcodes. Fastq files were generated with Illumina DRAGEN BCL Convert v3.8.4. Heatmaps were generated by Morpheus (Broad Institute).

The RNA-seq analysis was performed using OneStopRNAseq [50]. Briefly, FastQC and MultiQC were used for raw reads quality control and QoRTs for post-alignment quality control [51, 52]. Reads were aligned to the reference genome assembly mm10 with star_2.7.5a and annotated with gencode.vM25.primary_assembly [53, 54]. Aligned exon reads were counted toward gene expression with featureCounts_2.0.0 with default settings [55]. Differential expression (DE) analysis was performed with DESeq2_1.28.1 [56]. Significantly differentially expressed genes (DEGs) were filtered with the criteria FDR < 0.05 (|LFC|) > 0.585. Gene set enrichment analysis were performed with GSEA [57]. Heatmaps of TPM values were generated by Morpheus (Broad Institute).

### Adipocyte size quantification

Inguinal white adipose tissues were harvested, sectioned, and stained with hematoxylin and eosin. H&E stained tissues were imaged at 20X magnification and adipocyte size (area in μm^2^) was quantified using Adiposoft plugin in ImageJ software [58]. All of the adipocyte areas counted by Adiposoft were included in the analysis and are represented as relative frequencies.

### Lentiviral transduction

KPCY-PTHrP^OE^ cells were previously generated by lentiviral-mediated overexpression of *Pthlh* [21]. Briefly, KPCY cells were transduced with pLenti-C-Myc-DDK-P2A-Puro Lentiviral Particles with EV (Origene PS100092) or Pthlh ORF (Origene MR201519L3) and stably selected with puromycin (8 µg/ml). Stable KPCY-PTHrP^OE^ cells were maintained with puromycin after selection.

### Statistical analysis

All data points are biological replicates and represent individual mice, tissues, or cell samples. The data was first tested for normal distribution, followed by the appropriate statistical test as indicated in the figure legends. Data are represented as mean ± standard deviation; \**P<0.05*; non-significant *P* values are indicated in the figure text or legend. All statistical analysis was performed using GraphPad Prism.

### Data and code availability

All sequencing data have been deposited in the Gene Expression Omnibus under the series GSE298308.

## ACKNOWLEDGEMENTS

This work was supported by an American Gastroenterology Association Bern Schwartz Research Scholar Award in Pancreatic Cancer (JRP); National Institute of Health (NIH) K99-R00 CA252153 (JRP); US Department of Defense (DOD) Pancreatic Cancer Research Program (PCARP) Idea Development Award HT9425-24-1-0784 (JRP and DG); Hopper-Belmont Inspiration Award (NB); Pancreatic Cancer Alliance (PCA) Postdoctoral Fellowship (NB); NIH Innate Immunity Training Program T32 AI095213 (JP and CJ); NIH Institute for Maximizing Student Development T32 GM135751 (CJ); Canadian Institutes of Health Research (CIHR) MOP142287 (RK); Pancreatic Cancer Detection Consortium, U01CA210240 (MAH); NCI Cancer Center Support Grant, P30CA36727; NCI Research Specialist, R50CA211462; NIH DK130852 (MPC); Breast Cancer Alliance Young Investigator Grant (EVW); Massachusetts Life Sciences Center Bits to Bytes grant (to Drs. Dori Schafter and Christina Baer); and Massachusetts Life Sciences Center Research Infrastructure grant (to Drs. Kate Fitzgerald and Christina Baer). We acknowledge and thank the Sanderson Center for Optical Experimentation (SCOPE; Dr. Christina Baer and Jill McConnell) and Morphology core facilities at the University of Massachusetts Chan Medical School; and Pancreatic Cancer Mouse Hospital and the Molecular Pathology and Imaging Core at the University of Pennsylvania.

