## Supplemental Figure Legends.pdf for "Pancreatic cancer cachexia is mediated by PTHrP-driven disruption of adipose *de novo* lipogenesis"

**Supplemental Figure 1.**
**(A)** Schematic depicting KPCY-Pthlh$^{HET}$ mouse alleles. Created in BioRender. Pitarresi, J. (2025) https://BioRender.com/gjhr3fw. **(B-E)** Quantification of tumor burden (B) and normalized tissue masses for iWAT (C), eWAT (D), and quad (E) from a subset of animals with matched primary tumor burden. qPCR **(F)** qPCR for *Trim63* (MURF1) and *Fbxo32* (ATROGIN1) expression in quad muscles from the indicated genotypes in the age-matched cohort. **(G)** Quantification of adipocyte area in iWAT from the indicated genotypes. **(H-I)** Adipose tissue (H) or muscle (I) mass versus tumor mass, indicating no correlation between tumor burden and tissue wasting. **(J)** Kaplan-Meier overall survival plots of pancreatic cancer patients stratified by high *PTHLH* (red) or low *PTHLH* (blue) expression from TCGA-PAAD database. Logrank P = $6.0 \times 10^{-3}$. Statistical analysis by Student unpaired *t* test (B, C, D, E, and F) or Mann–Whitney *U* test (C), with significance indicated (*, $P < 0.05$). Error bars indicate standard deviation (SD).

**Supplemental Figure 2.**
**(A)** *Pthlh* expression in a panel of KPCY cell lines with the indicated non-cachectic clone (blue) used for subsequent studies. **(B)** *Pthlh* expression after lentiviral-mediated overexpression (PTHrP$^{OE}$) in the non-cachectic KPCY clone from (A). **(C)** WST-1 cell viability assay measuring optical density (OD) in the indicated genotypes. **(D)** Quantification of adipocyte area in iWAT from the indicated genotypes. **(E)** qPCR for *Trim63* (MURF1) and *Fbxo32* (ATROGIN1) expression in quad muscles from the indicated genotypes. **(F)** Quantification of endpoint tumor burden in KPCY-EV or KPCY-PTHrP$^{OE}$ tumor bearing mice. **(G)** Quantification of tumor burden in the primary tumor sized matched cohort of KPCY-EV or KPCY- PTHrP$^{OE}$ tumor bearing mice. **(H-I)** iWAT (H) and quad (I) masses in the primary tumor size matched cohort of KPCY EV or KPCY-PTHrP$^{OE}$ tumor bearing mice in (G). Statistical analysis by Student unpaired *t* test (B, E, F, G, H, I), ANOVA (C) or Mann–Whitney *U* test (D), with significance indicated (*, $P < 0.05$). Error bars indicate standard deviation (SD).

**Supplemental Figure 3.**
**(A)** Human *PTH1R* expression from GTEX database in all tissues analyzed. Adipose tissues are indicated with a red arrow and muscle tissues with a green arrow. **(B)** Single-cell RNA-seq clustering plots (left) and *Pth1r* expression (right) from the Emont *et al.*, murine adipocyte single cell atlas [25]. **(C)** Quantification of adipocyte area in iWAT from the indicated genotypes. **(D)** qPCR for *Trim63* (MURF1) and *Fbxo32* (ATROGIN1) expression in quad muscles from the indicated genotypes. Statistical analysis by Mann–Whitney *U* test (C) or Student unpaired *t* test (D), with significance indicated (*, $P < 0.05$). Error bars indicate standard deviation (SD).

**Supplemental Figure 4.**
**(A-B)** Heatmap of differentially expressed genes (A) and Principal Component Analysis (B) from RNA-seq of iWAT from KPCY-EV and KPCY- PTHrP$^{OE}$ tumor bearing mice. (C)

Heatmap of the top 50 up- and down-regulated genes in the iWAT from KPCY-EV versus KPCY- PTHrP$^{OE}$ mice.

**Supplemental Figure 5.**
**(A)** Western blot analysis of FASN and ACTIN in iWAT from KPCY-EV orthotopic tumor-bearing WT or Fasn$^{Adipo\text{-}cKO}$ hosts. **(B)** Quantification of adipocyte area in iWAT from the indicated genotypes. **(C-D)** Normalized tumor mass (C) and quad muscle mass (D) in KPCY-EV orthotopic tumor-bearing WT or Fasn$^{Adipo\text{-}cKO}$ hosts. **(E)** Representative H&E images of quads from KPCY-EV orthotopic tumor-bearing WT or Fasn$^{Adipo\text{-}cKO}$ hosts. Scale bars, 100 µm. **(F-G)** Normalized iWAT mass (F) and eWAT mass (D) in non tumor bearing WT or Fasn$^{Adipo\text{-}cKO}$ hosts. Statistical analysis by Mann–Whitney *U* test (B) or Student unpaired *t* test (C, D, F, and G), with significance indicated (*, $P < 0.05$). Error bars indicate standard deviation (SD).

**Supplemental Figure 6.**
**(A)** Quantification of adipocyte area in iWAT from the indicated GEMM genotypes. **(B)** qPCR for *Trim63* (MURF1) and *Fbxo32* (ATROGIN1) expression in quad muscles from the indicated GEMM genotypes. **(C)** Quantification of adipocyte area in iWAT from mice orthotopically implanted with tumor cells of the indicated genotypes. **(D)** qPCR for *Trim63* (MURF1) and *Fbxo32* (ATROGIN1) expression in quad muscles from mice orthotopically implanted with tumor cells of the indicated genotypes. **(E)** qPCR of *Acly, Acaca*, and *Acss2* in iWAT from KPCY orthotopic tumor-bearing hosts treated with anti-IgG or anti-PTHrP. Statistical analysis by Mann–Whitney *U* test (A and C) or Student unpaired *t* test (B, D, and E), with significance indicated (*, $P < 0.05$). Error bars indicate standard deviation (SD).
