## Supplementary figures and images for "Pancreatic cancer cachexia is mediated by PTHrP-driven disruption of adipose *de novo* lipogenesis"

### Supplemental Figure 1

#### SUPPLEMENTAL FIGURE 1

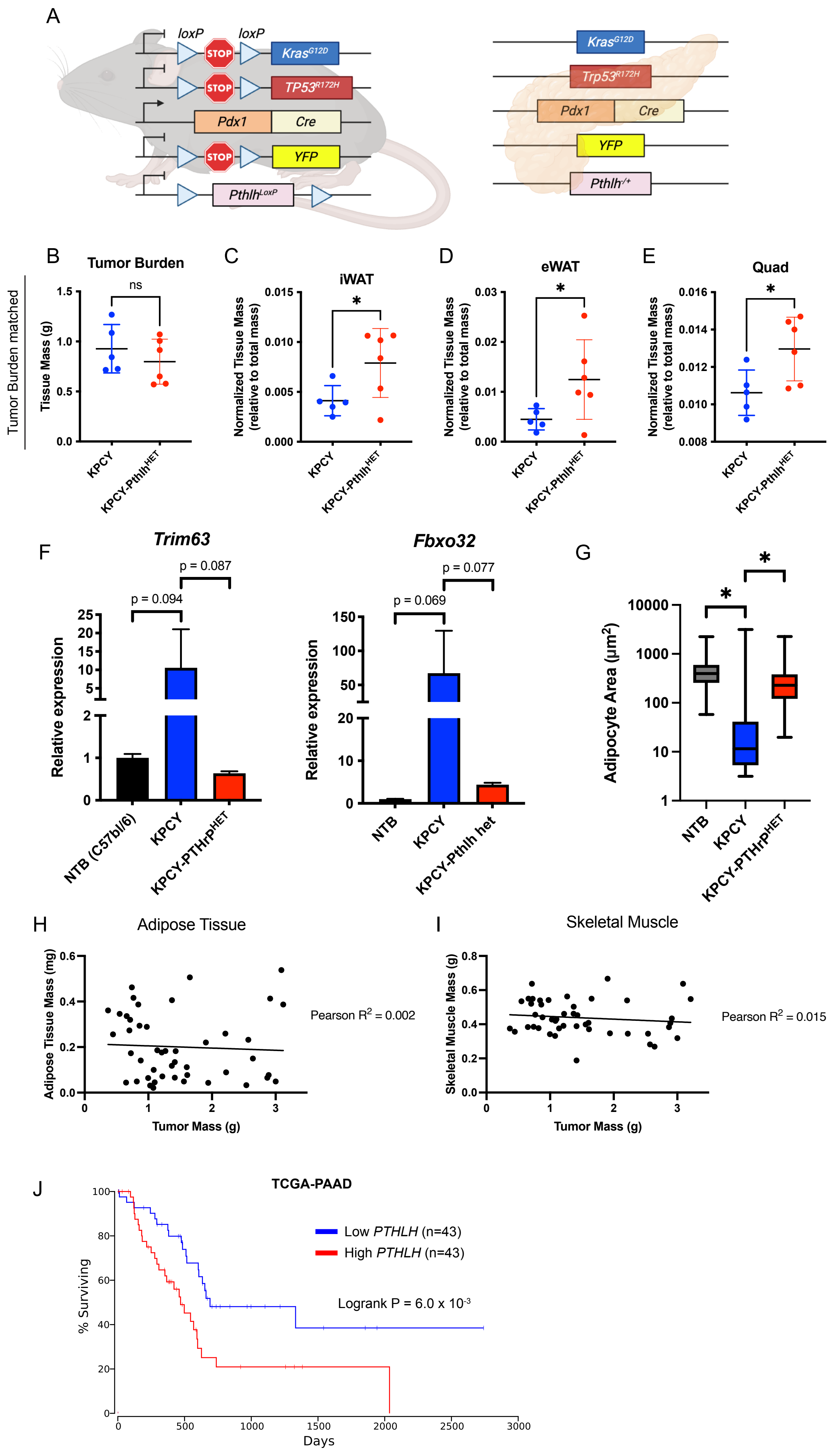

### Supplemental Figure 2

#### SUPPLEMENTAL FIGURE 2

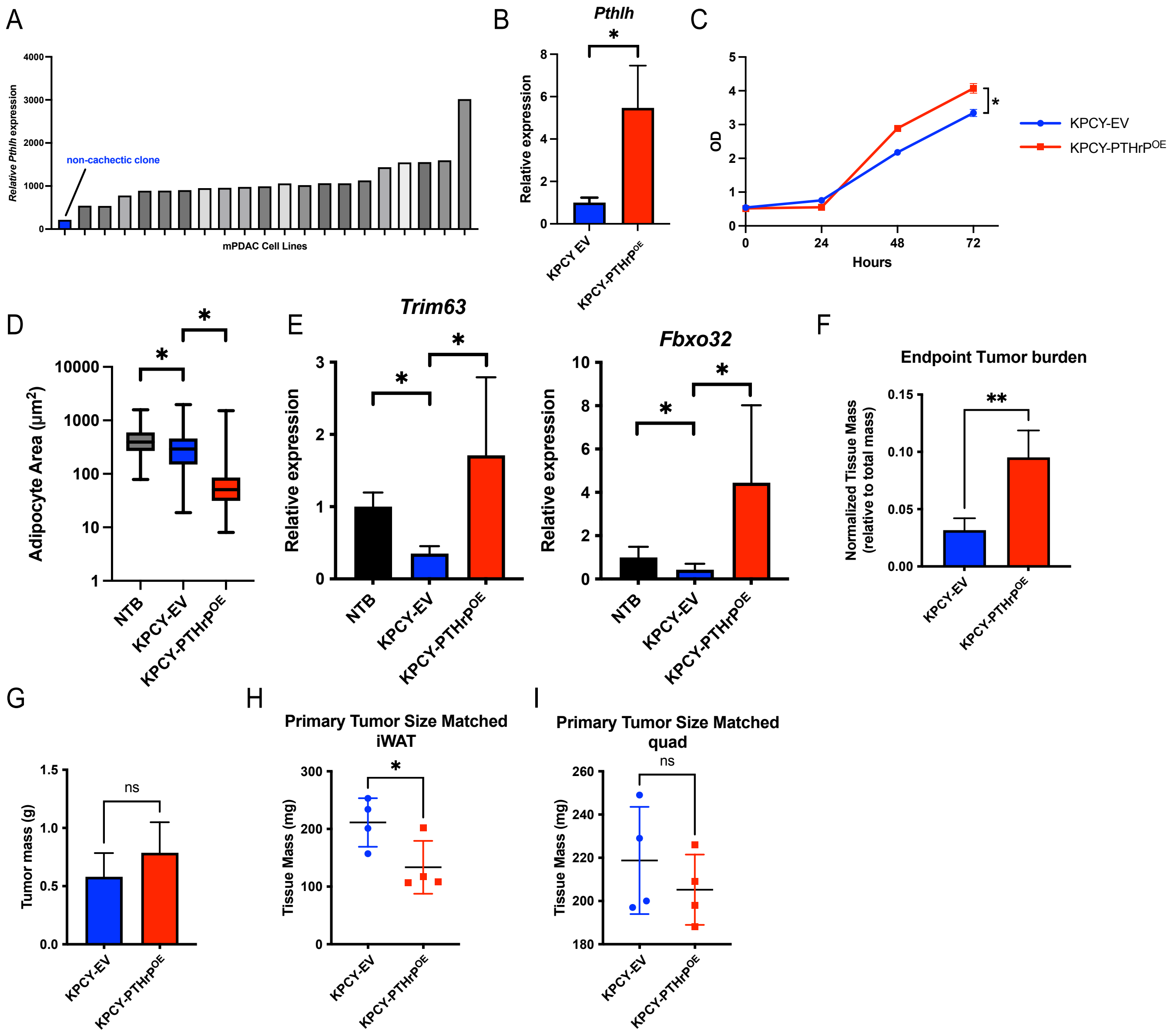

### Supplemental Figure 3

#### SUPPLEMENTAL FIGURE 3

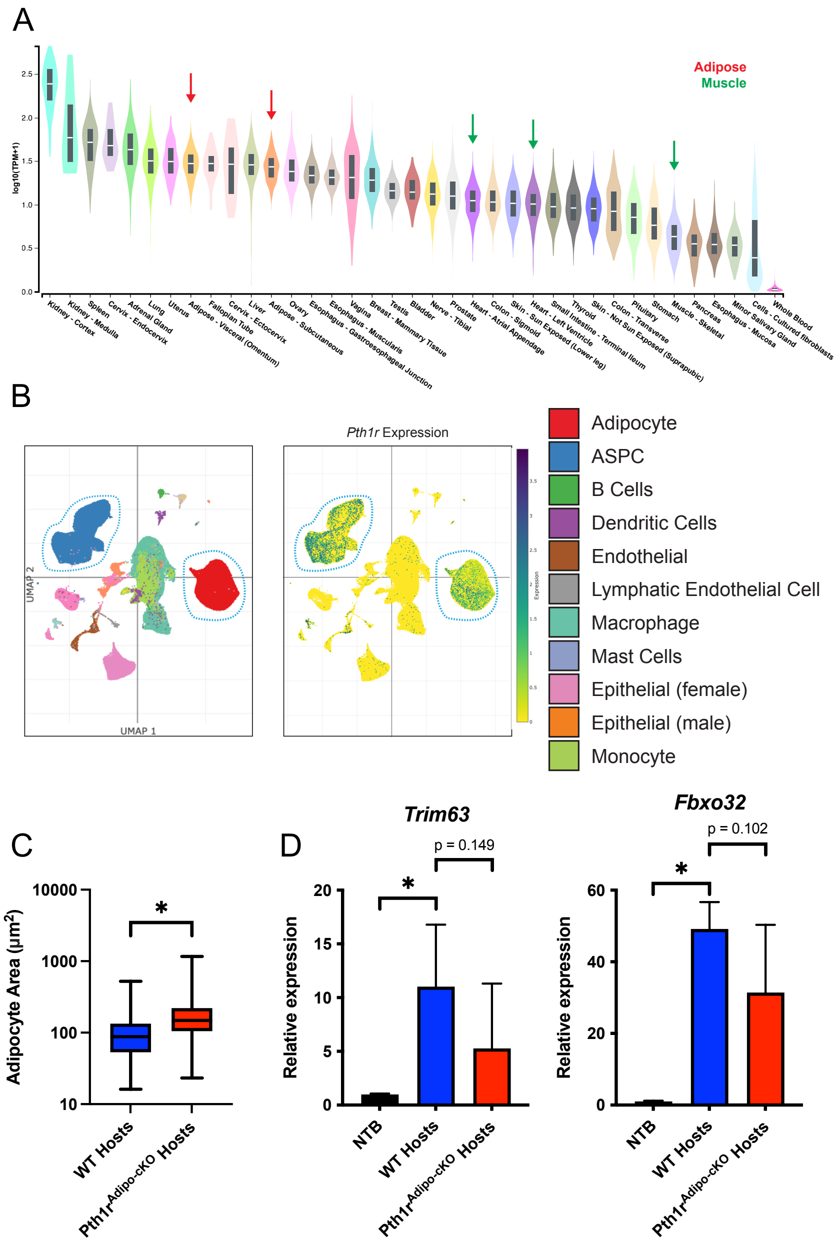

### Supplemental Figure 4

#### SUPPLEMENTAL FIGURE 4

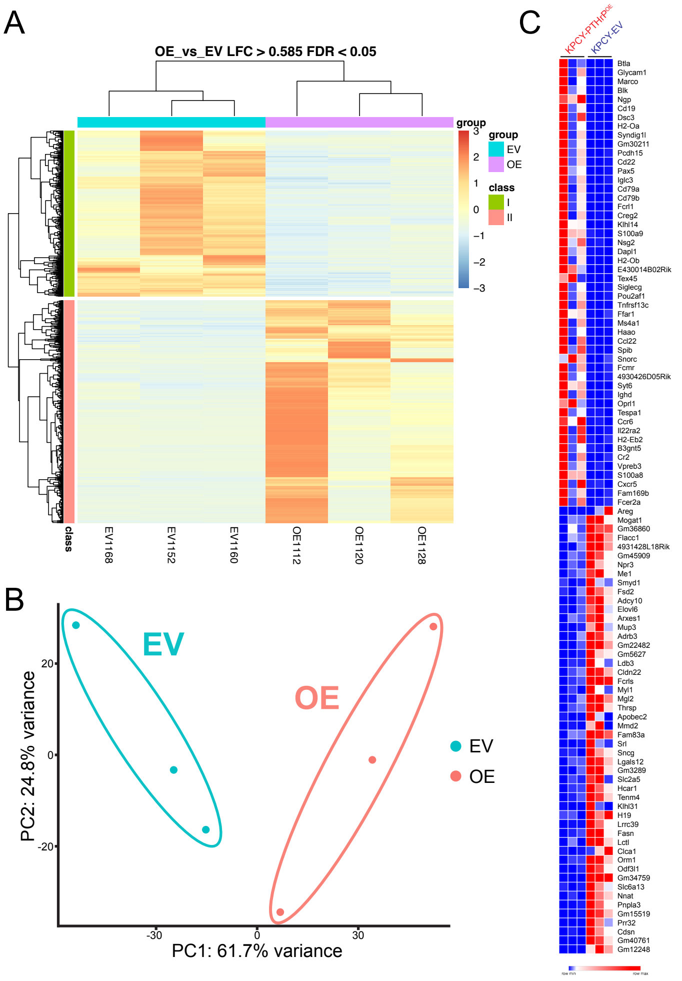

### Supplemental Figure 5

#### SUPPLEMENTAL FIGURE 5

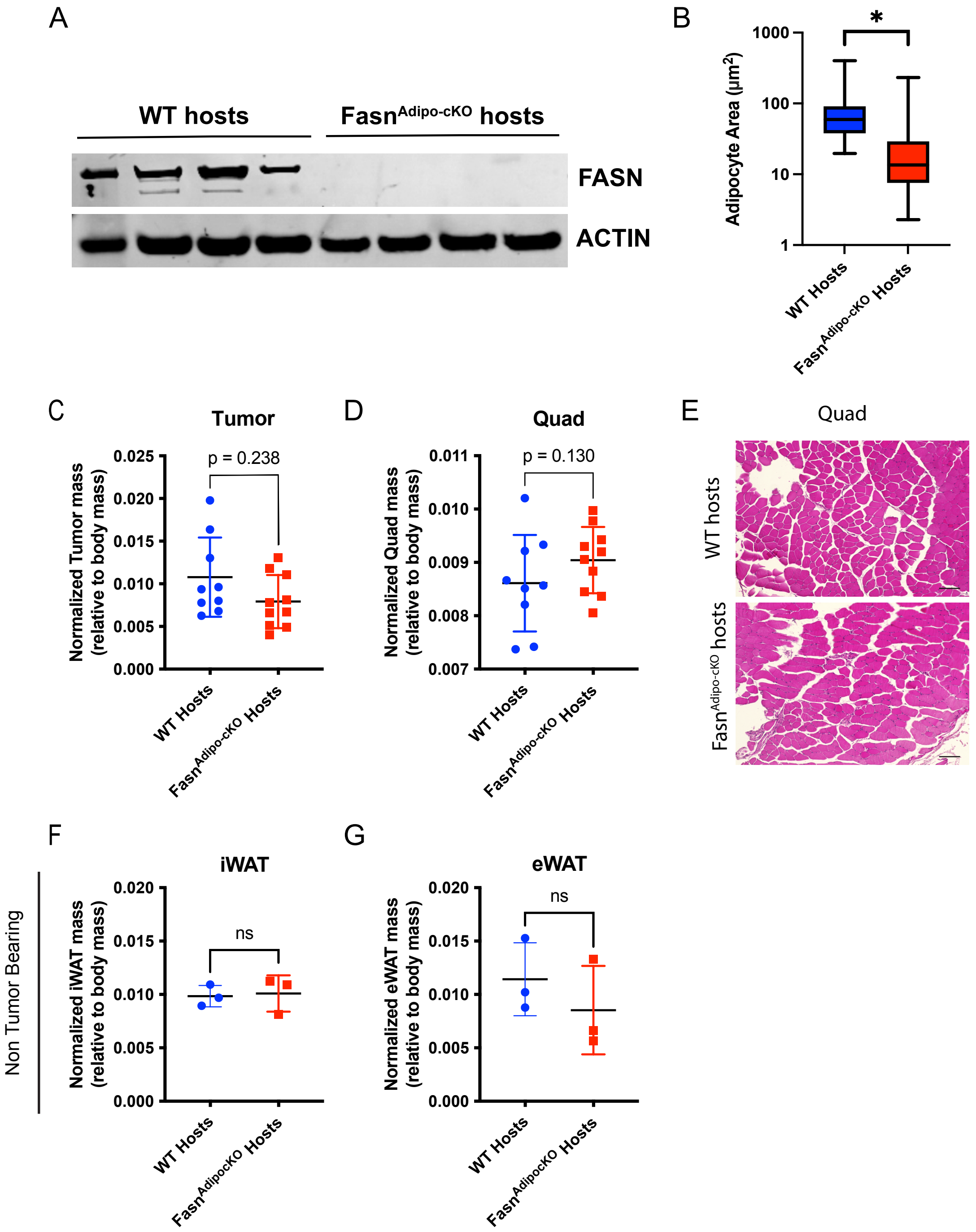

### Supplemental Figure 6

SUPPLEMENTAL FIGURE 6

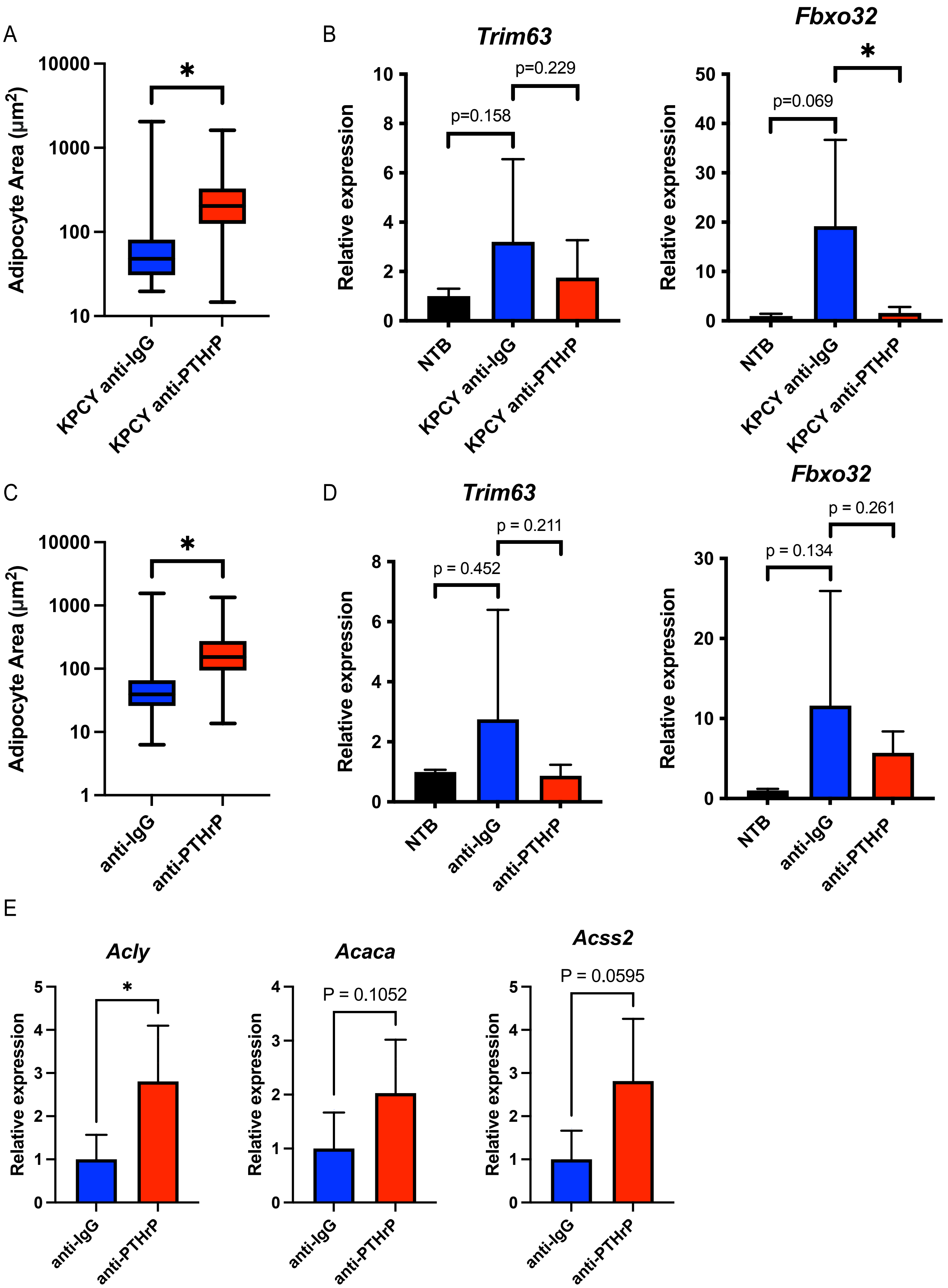
